# Chip (Ldb1) is a new cofactor of Zelda forming a functional bridge to CBP during zygotic genome activation

**DOI:** 10.1101/2025.04.24.650404

**Authors:** Charalampos Chrysovalantis Galouzis, Yacine Kherdjemil, Mattia Forneris, Rebecca R. Viales, Raquel Marco-Ferreres, Eileen E.M. Furlong

## Abstract

The cofactor Ldb1 (Chip) is linked to many processes in gene regulation, including enhancer-promoter communication, inter-chromosomal interactions and enhanceosome-cofactor-like activity. However, its functional requirement and molecular role during embryogenesis has not been assessed to date. Here, we used optogenetics (iLEXY), to rapidly deplete *Drosophila* Ldb1 (Chip) from the nucleus at precise time windows. Remarkably, this pinpointed the essential window of Chip’s function in just one-hour of embryogenesis, overlapping zygotic genome activation (ZGA). We show that Zelda, a pioneer factor essential for ZGA, recruits Chip to chromatin, and both factors regulate concordant changes in gene expression, suggesting that Chip is a cofactor of Zelda. Surprisingly, Chip is not required for chromatin architecture at these stages, instead it recruits CBP, and is essential for the placement of H3K27ac. Taken together, our results identify Chip (Ldb1) as a functional bridge between Zelda and the coactivator CBP to regulate gene expression in early embryogenesis.

## INTRODUCTION

Transcription factors (TFs) regulate gene expression through the recruitment of cofactors^1^. By definition, cofactors cannot bind to DNA directly, but are recruited to DNA by TFs or chromatin modifications, and have diverse functions, including ATPase dependent chromatin remodelers such as SWI/SNF^2^, chromatin modifying enzymes such as CBP/p300^3^, and components of the Mediator complex^4,5^. Most cofactors bind to thousands of regions across the genome, and therefore have a widespread impact on transcription, in addition to having diverse roles in the process of gene regulation^6,7^. CBP/p300, for example, catalyzes the acetylation of histones as well as other proteins, including TFs and components of the RNA polymerase machinery^3,8,9^. Other cofactors, such as the Mediator complex, act as a scaffold in large protein complexes, and are essential for RNA Pol II recruitment^5,10^. Cofactors therefore relay information between TFs bound to an enhancer and the basal transcriptional machinery at the promoter, and can either activate or repress transcription.

Ldb1 (called Chip in *Drosophila*) was the first cofactor identified to be essential for long-range enhancer-promoter (E-P) communication. It was discovered in a genetic screen in *Drosophila*, looking for genes that are essential for the *wing margin* enhancer to activate the promoter of its target gene, *cut*, ∼80kb away^11,12^. Since then, the role of its mammalian homolog, Ldb1, in E-P looping has been demonstrated at the *β-globin* locus in erythroid cells^13,14^, and is even sufficient to form a loop in both mouse^15^ and human^16^ cells. Similarly, during mammalian olfactory sensory neuron differentiation, Ldb1 can bridge multiple enhancers in both *cis* and *trans* to the promoter of the single olfactory receptor allele that is expressed^17^. In line with other cofactors, Chip (Ldb1) appears to be highly pleiotropic, with other functions in the regulation of gene expression, including being a component of the Wnt enhanceosome, suggested to prime enhancers for Wnt responses in *Drosophila* S2 cells^18^. This is likely a conserved function as Ldb1 is also associated with Wnt signaling in mice^19^. Ldb1/Chip is also a component of several conserved complexes from *Drosophila* to mammals^20–23^ including its association with Pannier/GATA-1, LMO, SSDP and members of the bHLH family that regulate sensory bristle differentiation in *Drosophila*^24^ and erythroid gene expression in mammals^25–28^. Small changes in the components of these complexes can lead to large differences in gene regulation, such as the replacement of GATA-1 by GATA-2 in the regulation of haematopoietic stem cell gene expression^19,29^. In the context of the *β-globin* gene cluster, Ldb1 bridges multiple events required for efficient E-P communication such as gene positioning, looping as well as gene activation through enriching and stabilizing transcriptional coregulators^13,14,30,31^. Despite these important functions, there has been very little investigation of the role of Ldb1 (Chip) during embryonic development in any organism to date.

This is largely due to the difficulty in studying the function of cofactors in embryos using traditional genetic tools such as knock-outs and knock-downs. Their pleotropic functions and very broad roles in transcription lead to very severe defects that are often lethal at very early stages of embryogenesis^32,33^. Much of our current knowledge therefore comes from careful biochemical studies, genetic dissection in yeast and cell culture models, with little genetic dissection in embryos. To overcome these limitations, we took advantage of the recently optimized iLEXY optogenetic system, which triggers extremely rapid nuclear export of proteins upon blue light exposure in the order of seconds. When the blue light is switched off, the protein of interest can return to the nucleus, enabling reversible, tightly-controlled temporal perturbations in living cells or embryos^34,35^.

Here we tagged the endogenous *chip* gene with the iLEXYs module to investigate its function for the first time during multiple stages of *Drosophila* embryogenesis. We focused on the first two-thirds of embryogenesis, which span the major events of zygotic genome activation (ZGA), gastrulation, the specification of major cell lineages, and the initiation of terminal differentiation for many tissues. We demonstrate that the iLEXY system can (i) efficiently deplete Chip (Ldb1) from chromatin, (ii) recapitulate the known segmentation phenotype in early embryos, and (iii) mimic the embryonic lethality of a loss-of-function mutant when depleted over all of embryogenesis. By performing fine-grained temporal perturbations, we could narrow down the most sensitive time period to Chip (Ldb1) depletion over the entirety of embryogenesis to just a one-hour window that coincides with ZGA. Using quantitative CUT&Tag, we show that Chip is a cofactor of the pioneer factor Zelda. Zelda is essential to recruit Chip to DNA, such that the binding and transcriptional responses of both factors strongly overlap after depletion of either Chip or Zelda in early embryos. Chip is not required for Zelda’s pioneering role to open chromatin, but is required to recruit the coactivator CBP, which subsequently deposits H3K27ac at specific regions of the genome during these early stages of embryogenesis. Surprisingly, Chip does not appear to have a role in E-P chromatin looping at this stage of embryogenesis. In contrast, the majority (if not all) of its functions are mediated through Zelda. This highlights a new role of Chip as a missing cofactor of Zelda, which acts as functional bridge to the coactivator CBP, leading to H3K27ac deposition at both enhancers and promoters during ZGA.

## RESULTS

### Chip (Ldb1) binding at enhancers and promoters is dynamic and correlated with transcription

As the role of Chip (Ldb1) has not been examined during embryogenesis to date, we first assessed its occupancy at four different time windows covering early to mid-embryogenesis, spanning the stages of ZGA and gastrulation (2-3h (stage 5), 2-4h (stages 5 to 8), to cell fate specification of major lineages (6-8h, stage 11) and the initiation of terminal differentiation (10-12h, stage 13). This identified 15,102 significant peaks bound by Chip at one or more time-point (Table S1, Methods), indicating broad binding across the genome. Almost half (48.4%) of the bound regions are promoter proximal (within 500bp of an annotated transcriptional start site (TSS)) (Fig. 1A, Fig. S1A, B), and are enriched for genes with development, metabolic and signaling functions (Fig. S1C-F). Roughly a third of peaks (5,236) are shared between 2-3h, 2-4h and 6-8h, while the majority are time-point specific, including 5,018 peaks at 6-8h and 563 peaks at 2-3h and 2-4h (Fig. 1B-D), suggesting alternative roles for Chip (Ldb1) across embryogenesis. Chip binding changes dramatically at 10-12h compared to the earlier time-windows, with a decrease in both the number (1,957) and quantitative signal of peaks (Fig. 1C), perhaps reflecting a reduced requirement at later stages. The distribution of binding also changes at 10-12h to predominantly distal regions (65.8% of peaks) (Fig. 1A). As Chip doesn’t have a DNA-binding domain, it is recruited to DNA indirectly by transcription factors (TFs). Searching for TF motifs in the time-point specific Chip peaks revealed different likely cofactors at each stage (Fig. 1D, Table S2), supporting Chip having different potential roles during embryogenesis. For example, GATAe and eBox motifs (Amos, Twist) are enriched at 6-8h, while a homeodomain motif (dve, oc) is enriched at 2-4h (Fig. 1D).

**Figure 1:**
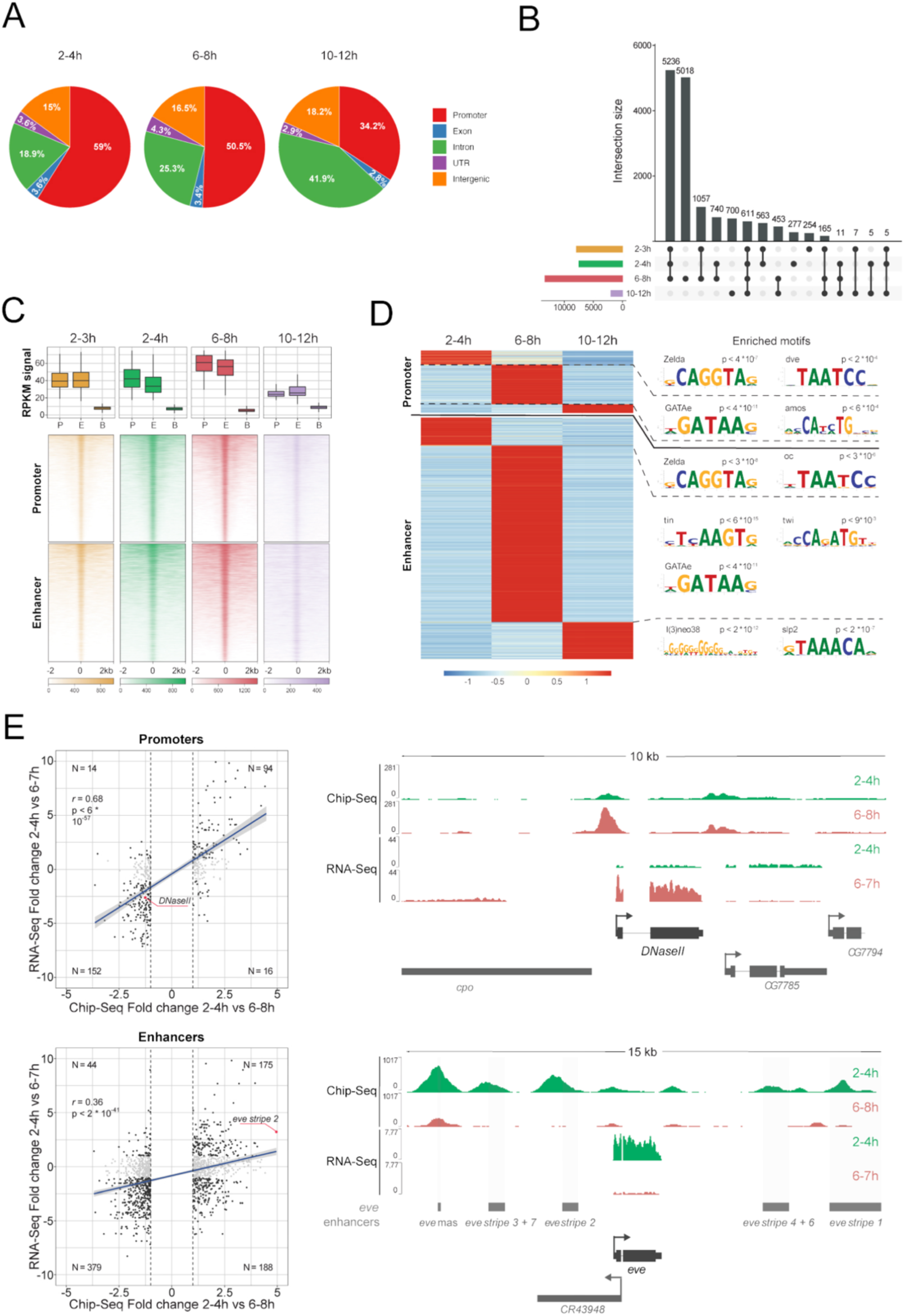
Chip binding to enhancers and promoters changes over embryogenesis. (**A**) Pie charts showing the percentage of overlap of Chip occupancy (ChIP-seq peaks) with genomic features at three different time windows of embryogenesis (2-4h, 6-8h and 10-12h). (**B**) Upset plot showing overlap of Chip peaks at four different time windows of embryogenesis (2-3h, 2-4h, 6-8h and 10-12h). (**C**) *Upper*: Boxplots of Chip ChIP-seq signal (RPKM) on enhancers (E), promoters (P) and background (B) (defined as inactive chromatin from DNase Hypersensitivity) at four time windows of embryogenesis (2-3h, 2-4h, 6-8h and 10-12h). *Lower*: corresponding heatmap of the signal, centered on the summit of enhancers or promoters. (**D**) Heatmap of time-window-specific Chip peaks at enhancers and promoters. Color scale represents the z-score normalized ChIP-seq signal across time-windows (rows). The most significant enriched motifs in those peaks (compared to all peaks) are depicted along with the adjusted p-value^90^. (**E**) Left: Scatter plots of the fold change of gene expression (RNA-seq) and the corresponding ChIP-seq signal at promoters (top left) and enhancers (bottom left) at 2-4h versus 6-7h. N = number of observations in each quadrant, *r* = Pearson correlation and p = p value. *Top right*: Genome tracks showing a gain of Chip occupancy at the promoter of *DNaseII* at 6-8h with in parallel gain of RNA-seq signal at 6-7h. *Bottom right*: Genome tracks showing the loss of Chip binding at *eve* enhancers at 6-8h, which is mirrored by a loss of RNA expression at 6-7h.

The changes in Chip binding across different stages of embryogenesis correlate with changes in gene expression – i.e. differential Chip binding between 2-4h and 6-8h at promoters and enhancers is globally correlated with differential gene expression between 2-4h and 6-7h embryos (*r*=0.68 and 0.36, respectively) (Fig. 1E, left). Chip binding, for example, at the promoter of the *DNaseII* gene increases from 2-4h to 6-8h, matching the activation of the gene’s expression at 6-7h (Fig. 1E, right). Similarly, Chip is bound to all five *eve* enhancers at 2-4h, matching the high levels of the gene’s expression at this stage, while the binding is strongly reduced at 6-8h when the gene’s expression is strongly decreased (Fig. 1E, right).

To explore the relationship between Chip binding and enhancer activity, we used a database of enhancers with characterized activity in transgenic embryos^36^, and examined the activity status of 933 enhancers bound by Chip at 2-4h and/or 6-8h of embryogenesis. This revealed significantly higher levels of Chip binding (quantitative signal) at 2-4h at enhancers that are active at 2-4h, compared to inactive enhancers at this stage of embryogenesis (p < 2.9e-54). Conversely, Chip binding is lower on these same enhancers at 6-8h when they are inactive (p < 4.6e-23) (Fig. S2A,B). The same trend holds true for enhancers active at 6-8h (and inactive at 2-4h) – ChIP binding is higher at 6-8h (Fig. S2C,D). Taken together, these results suggest a widespread and dynamic role for Chip in enhancer activation and the regulation of gene expression during embryogenesis, which we directly assess below.

### Optogenetic nuclear depletion of Chip (Ldb1) phenocopies *chip* loss-of-function mutant embryos

To better understand the function of Chip (Lbd1) during embryonic development in *Drosophila*, we took advantage of our recently optimized optogenetic approach (iLEXY) to very rapidly and reversibly deplete Chip from the nucleus^34,35^. As iLEXY functions at the protein level, it depletes both maternally deposited and zygotically expressed protein from the nucleus, which is crucial for maternally deposited genes like Chip. Using CRISPR-Cas9 mediated homology directed repair, we tagged the endogenous *chip* gene at the C-terminal with the slow cycling iLEXYs variant^35^, hereafter Chip::iLEXYs (Fig. 2A). The *chip::iLEXYs* line is homozygous viable and fertile when kept under safe light (which we refer to as ‘Dark’), and the fusion protein localizes to the nucleus as expected^12^ (Fig. 2B and S3A), indicating that the iLEXYs tag does not perturb the function of Chip.

**Figure 2:**
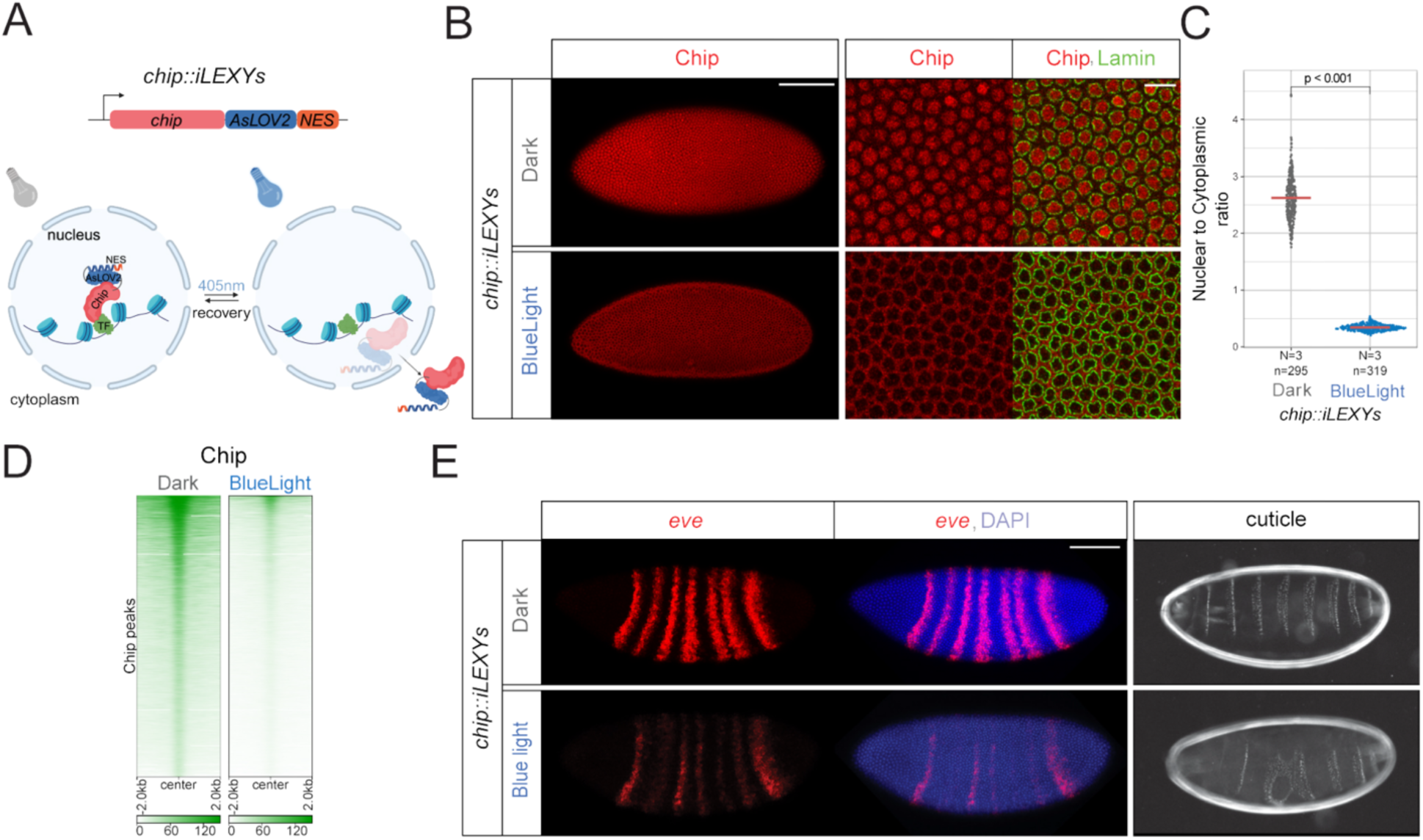
iLEXY efficiently depletes Chip from chromatin and phenocopies loss-of-function mutant embryos. (**A**) Schematic representation of the iLEXY system used for the modulation of Chip localization. The endogenous *chip* gene was tagged with iLEXY, which upon blue light exposure leads to unfolding of the LOV domain and exposure of a nuclear export signal (NES). When the blue light is switched off, the protein returns to the nucleus. (**B**) Immunostaining of Chip (red) and Lamin (green) stage 5 embryos (2-4h) in the dark and after blue light exposure (left) and zoom in (right). Scale bar = 100 μm (left) and, 10 μm (right). (**C**) Quantification of the fluorescence intensity of Chip in nuclei compared to cytoplasm. Horizontal lines = median, N = number of embryos, n, number of nuclei, p-value from t-test. (**D**) Heatmap of Chip CUT&Tag normalized signal (using spike in) in 2-4h *chip::iLEXYs* embryos in dark and blue light exposure. Peaks are centered on the peak summits. (**E**) *Left*: RNA FISH for *eve* (red) (left) and DAPI (blue) in stage 6 embryos in dark and after blue light exposure. Scale bar = 100 μm (top right) for RNA FISH embryos. *Right*: cuticular preparation of stage 17 embryos in dark and blue light exposure. Schematic was created with Biorender.com.

To determine if Chip can be depleted from the nucleus, we incubated homozygous embryos in blue light, using a programmable LED blue light box^35^, under conditions that do not perturb the viability of control (*yw*) embryos. After blue light exposure, embryos showed a strong nuclear depletion of Chip and relocalization to the cytoplasm, as seen by immunostaining of stage 5 (2-3hr) embryos (Fig. 2B). Quantification of the nuclear to cytoplasmic ratio in dark versus blue light incubated *chip::iLEXYs* 2-4h embryos, showed a 10-fold signal reduction (Fig. 2C). Moreover, Chip is strongly depleted from chromatin after blue light exposure, as measured using spike-in CUT&Tag in nuclei from dark versus blue light embryos (Fig. 2D, S5A). For quantification, nuclei from an evolutionary distant *Drosophila* species (*D. virilis*) were used for the spike-in control for all experiments (Methods).

Importantly, optogenetic nuclear depletion is sufficient to recapitulate the *chip* loss-of-function mutant phenotype. Specifically, *in-situ* hybridisaton of *eve* showed a similar misexpression after iLEXY-mediated nuclear depletion as observed in germline clones using the *chip^e5.5^* loss-of-function allele^12^ (Fig. 2E). All seven *eve* stripes are affected to varying degrees, although stripes 1 and 7 to a lesser extent as also observed in *chip^e5.5^* germline clone embryos (Fig. 2E). Consistent with the segmentation phenotype of *chip^e5.5^* germline clones, cuticles of the *chip::iLEXY* embryos showed segmentation defects after blue light exposure, with irregular and fused denticles along the anterio-posterior axis (Fig. 2E, right).

Together, these results demonstrate that iLEXY-mediated nuclear depletion can very efficiently deplete Chip from chromatin and recapitulate the known *chip* loss-of-function phenotypes.

### Depletion of Chip (Ldb1) leads to diverse transcriptional changes at different stages of embryogenesis

To investigate how Chip regulates gene expression, we performed RNA-seq in *chip::iLEXYs* embryos at different time windows, which were either maintained in dark or exposed to blue light, leading to the nuclear depletion of Chip. This ensures that the embryonic stages are matched, in addition to the genotype. At the early time windows (2-3h and 2-4h), 448 and 415 gene are significantly upregulated (|log2 fold change| > 1 and FDR < 0.01) and 391 and 309 genes downregulated in their expression, respectively, in *chip::iLEXYs* embryos in blue light versus dark (Fig. 3A, B, E, Table S3). The genes that are deregulated in 2-3h embryos and bound by Chip at their promoter (Chip peak in ±500bp from TSS) have functions related to development as well as the regulation of transcription (Fig. S4A). The transcriptional changes observed at 2-3h and 2-4h are highly overlapping, as expected (Fig. S4B,C). The majority of deregulated genes, between the early (2-3h and 2-4h) and mid/late-stages (6-7h, 10-12h), are time point specific, in keeping with the dynamic binding of Chip, supporting Chip’s different functions at different stages of embryogenesis (Fig. S4B). While the transcriptional response between upregulated and downregulated genes is quite balanced at early stages, at mid-embryogenesis (6-7h) Chip depletion leads to a predominant upregulation of gene expression with 617 genes going up versus only 49 genes going down upon 1h blue light exposure (Fig. 3C, Fig. S4D), suggesting a more repressive role in gene expression at this stage. Interestingly, at 10-12h, very few genes were affected after 1h (11-12h) blue light exposure with only 3 downregulated and 66 upregulated genes (Fig. 3D), indicating a very limited role for Chip in gene expression at this time window in keeping with its reduced occupancy (Fig. 1C).

**Figure 3:**
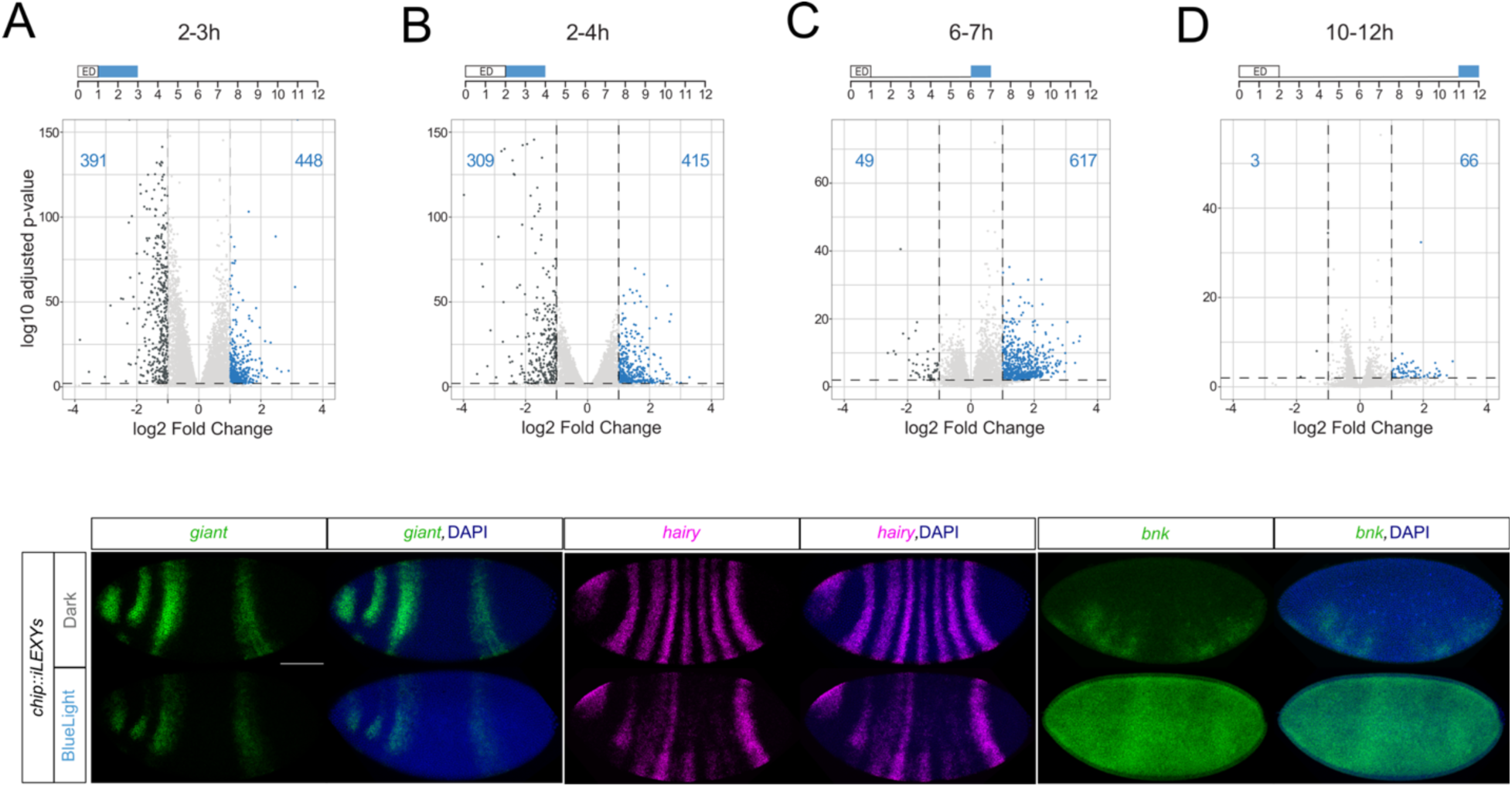
Different transcriptional responses to Chip depletion at different stages of embryogenesis. (**A-D**) Volcano plots showing the differentially expressed genes at different time windows of embryogenesis after Chip depletion: blue light exposure (indicated in blue) from1-3h sampled at 2-3h (**A**), 2-4h exposure sampled at 2-4h (**B**), 6-7h exposure sampled at 6-7h (**C**), and 11-12h exposure sampled at 10-12h (D). Blue = genes upregulated after Chip depletion, dark gray = genes downregulated, using |log2 fold change| > 1 and FDR < 0.01. ED = egg deposition. (**E**) RNA FISH of the downregulated genes *giant* (green) and *hairy* (magenta), and the upregulated gene, *bnk* (green), in dark (upper) and after blue light depletion (lower). Blue = DAPI. Scale bar = 100 μm.

We confirmed the misexpression of three new Chip target genes by *in-situ* hybridization, *giant* and *hairy*, which have decreased and *bottleneck* (*bnk*) with increased expression upon Chip nuclear depletion (Fig. 3E). We note that the transcriptional changes observed by whole embryo RNA-seq (Fig. 3A-D, S4C,D) are underestimates of the full transcriptional response to Chip depletion as they fail to capture variation in the spatial patterns of genes that remain partially expressed. For example, *eve* expression was not detected as significantly downregulated (using our stringent cut-offs) in the 2-3h RNA-seq experiment (having 20.2% lower expression in blue light, FDR < 4.3 × 10^−8^), despite a clear mis-regulation of the *eve* striped pattern in 2-4h embryos (Fig. 1D). Given this, and the changes we observed in *giant* and *hairy* expression, we examined the expression of a number of other segmentation genes by in-situ hybridization, and identified reduced expression for *hunchbac*k (*hb*) and *tailless* (*tll*) in *chip::iLEXYs* embryos at 2-4h upon blue light exposure, despite not being detected as significantly downregulated by RNA-seq (Fig. S4E).

### Time-controlled depletion of Chip (Ldb1) uncovers a narrow essential window for its function during embryogenesis

Removal of both maternal and zygotic Chip, in *chip^e5.5^* germline clones, is embryonic lethal^12^, while zygotic homozygous *chip* mutants die before the third-instar larval stage^11^. However, using these conventional genetic tools it is not possible to determine if Chip is required continuously during embryonic development or whether there is a specific time window in which it is necessary. Taking advantage of the speed and reversibility of the iLEXY system, we assessed the necessity of Chip for embryonic viability during successive time windows by quantifying embryo hatching rate into first-instar larvae. For all measurements, *chip::iLEXYs* embryos were incubated along-side control (*yw*) embryos to control for temperature and any potential effects of blue light exposure on viability. Importantly, *chip::iLEXYs* embryos incubated in the dark showed a similar viability rate compared to control embryos, demonstrating that the iLEXYs tag has no effect on viability (Fig. 4A). To mimic loss-of-function germ line and zygotic mutant embryos, we collected embryos for 30 minutes and then immediately exposed them to blue light for all of embryogenesis (24h in blue light). Blue light exposure (leading to Chip nuclear depletion) throughout all of embryogenesis is lethal (99% (529/532) lethality), while it has little or no effect on the control (*yw*) embryos (Fig. 4A). We next kept 30-minute embryo collections in the dark (allowing them to develop normally) for increasingly longer time windows at the beginning of embryogenesis, and then exposed them to blue light for the following time periods: 1h40’-24h, 3-24h and 6-24h (Fig. 4A). This revealed a very strong requirement of Chip during the first hours of embryogenesis, while the effects are less pronounced at later stages (Fig. 4A). Depletion from 1h40’ to the end of embryogenesis, for example, resulted in 93% (376/403) lethality, while depletion from 6h onwards resulted in only 40% (222/555) lethality.

**Figure 4:**
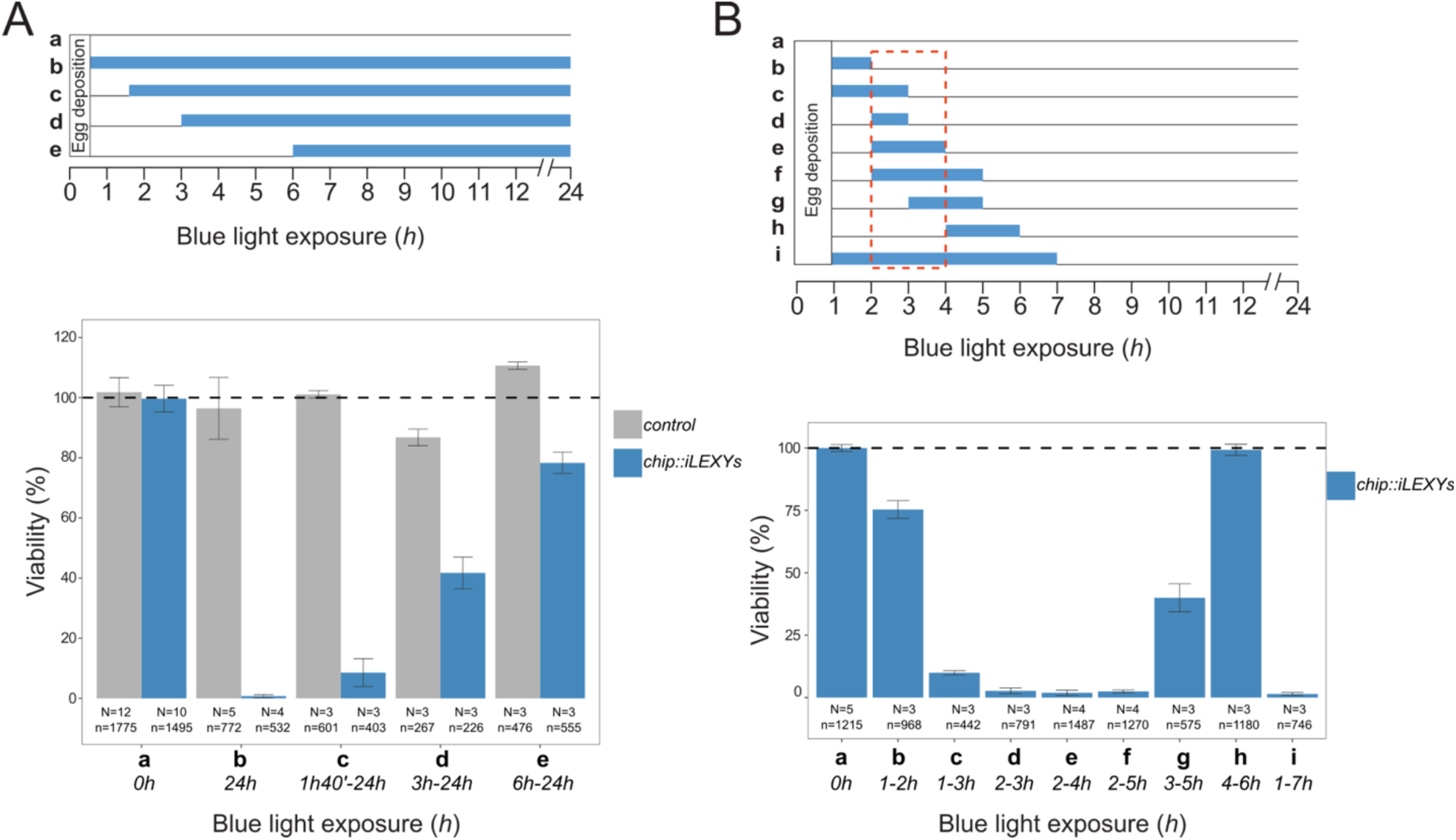
Chip is essential for embryonic viability in a narrow time window early (2-3h) in embryogenesis. (**A**) *Top*: Time windows showing the duration of blue light exposure (indicated in blue) after egg deposition. Conditions are indicated by letters (a-e). *Bottom*: Bar plots (mean ±SD) of embryonic viability (measured by hatching rate to first-instar larvae). Gray = control (*yw*) embryos, and blue = *chip::iLEXYs*. Percentages presented are normalized to the control embryos exposed to blue light in the same conditions (a). (**B**) *Top*: Short time windows of blue light exposure (indicated in blue) after egg deposition (ED). Conditions are indicated by letters (a-i). *Bottom*: Bar plots (mean ±SD) of embryonic viability of *chip::iLEXYs* embryos (hatching rate to first-instar larvae). Percentages are normalized to the *chip::iLEXYs* embryos not exposed to blue light (a). N = number of biological replicates, n = number of embryos.

To narrow down the essential time window, we depleted Chip for short time periods (3hr, 2hr and 1hr time windows). Such a high-temporal assessment of a gene’s function is not possible with classic genetic approaches. Depletion of Chip for the first third of embryogenesis (1-7h), and then allowing it to return to the nucleus (switching off the blue light) for the rest of embryogenesis is almost completely lethal, while depletion between 4-6h has almost no impact on viability (14% (170/1180) lethality) (Fig. 4B). This indicates that Chip is essential in the first four hours of embryogenesis. Applying two- and one-hour depletions, we narrowed down the essential window of Chip requirement to between 2 to 3h (or likely 3.5h, for full nuclear protein recovery) of early embryogenesis (Fig. 4B). The five-time windows overlapping a 2-3h depletion (Fig. 4B, conditions c, d, e, f and i) have only 8.5%, 2.3%, 1.6%, 2.1% and 1.2% viability, respectively. In contrast, Chip depletion in time windows before (condition b) and after (condition h) this critical early window has minimal impact, having 75% and 100% viability rate, respectively (Fig. 4B).

In summary, these results demonstrate that the nuclear depletion of Chip is embryonically lethal, consistent with the loss-of-function allele^12^ and that exposure to blue light successfully perturbs Chip protein function in the *chip::iLEXYs* line. Chip appears to be largely dispensable for viability in the latter two-thirds or more of embryogenesis, which is very surprising given its ubiquitous expression and many reported roles in the regulation of gene expression. However, it does fit with the reduced binding (Fig. 1C) and the minimal changes in gene expression (Fig. 3D) that we observed at 10-12 hours. Rather, we could pin-point a very narrow window of embryogenesis in which Chip is essential for viability, and interestingly, this critical window (2-3h) overlaps with the period of zygotic genome activation (ZGA).

### Zelda recruits Chip to sites where they are both required to regulate gene expression

Since Chip does not carry a DNA-binding domain, a TF must recruit it to chromatin. The most strongly enriched motif in the regions bound by Chip specifically at 2-4h of embryogenesis (overlapping the critical window of Chip function described above), is the Zelda motif, being highly enriched at both promoters (p < 3*10^-8^) and enhancers (p < 3*10^-7^) (Fig. 1D, Table S2). Zelda is a pioneer TF with a major role in ZGA during early stages of *Drosophila* embryonic development^37–39^. This suggests that Zelda might recruit Chip in this early phase of embryogenesis. To investigate this, we first compared the binding of Chip and Zelda, performing CUT&Tag for Chip in *chip::iLEXYs* (not exposed to blue light (dark)) and Zelda in control (*yw)* embryos at 2-4h. The binding profiles of Chip and Zelda are highly overlapping, with 86.6% (9,286/10,719) of Chip peaks being co-bound by Zelda (Fig. 5A, Table S4), with many of the remaining peaks having weaker (but consistent) Zelda enrichment, suggesting that they are subthreshold peaks (Fig. S5E).

**Figure 5:**
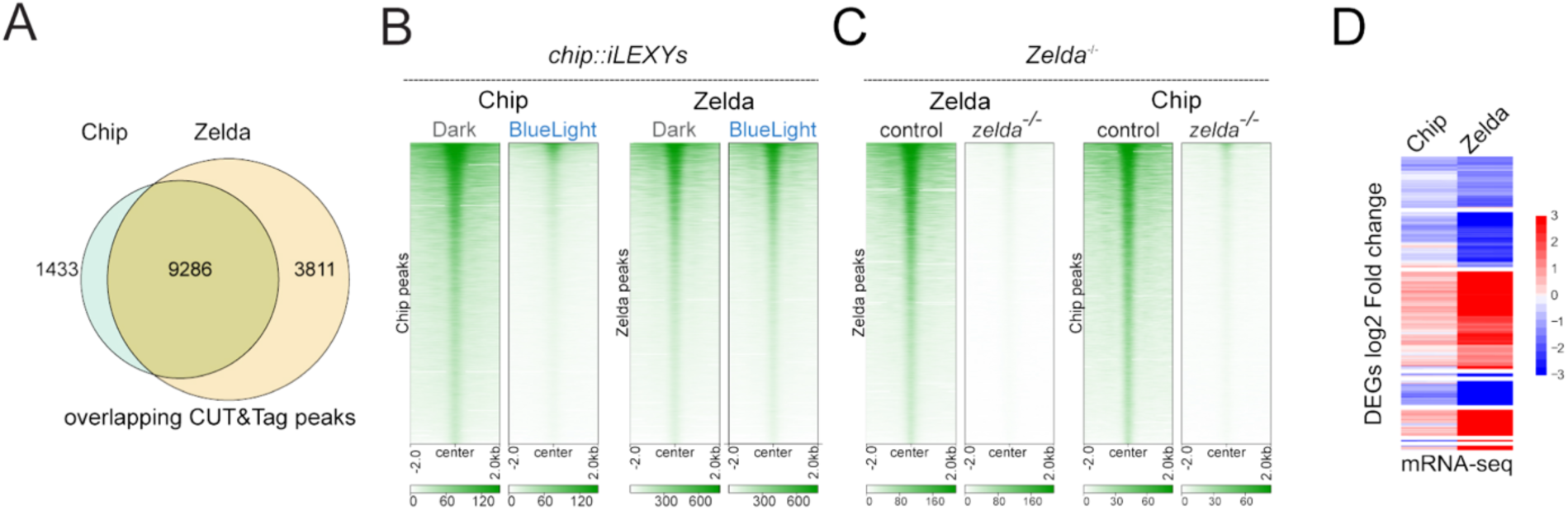
Zelda recruits Chip to chromatin and both genes have similar effects on gene expression. (**A**) Venn diagram showing overlap of Chip (CUT&Tag peaks in 2-4h *chip::iLEXYs* embryos in the dark) and Zelda (CUT&Tag peaks in 2-4h control *(yw)* embryos). (**B-C**) Heatmaps of Chip and Zelda CUT&Tag signal in 2-4h embryos, normalized using signal from spike-in *D. virilis* nuclei, in the following conditions: *chip::iLEXYs* embryos in the dark and after blue light exposure (Chip data is the same as in Fig. 2D) (**B**), or control and *zelda^-/-^* germ-line clone embryos (**C**), centered on the peak summits. (**D**) Heatmap of differentially expressed genes (DEGs), showing log2 fold-change of RNA-seq after Chip depletion at 2-3h, and in *zelda^-/-^* germline clone embryos during stage 5 (approx. 2-3h) (dataset from *Schulz et al*., 2015^40^).

We next determined if Zelda is required for Chip binding and vice versa, by quantitatively assessing the occupancy of Chip in *zelda* germ-line clone embryos (*zelda^-/-^*), and of Zelda in *chip::iLEXYs* 2-4h embryos after blue light deletion, using spike-in CUT&Tag (Methods). The occupancy of Chip in *chip::iLEXYs* embryos after blue light exposure and Zelda in *Zelda^-/-^* embryos are dramatically reduced, with 98.2% (10,527/10,719) and 96.2% (10,884/11,313) of peaks being significantly depleted [FDR < 0.05 and |log2FC| > 0.5]), respectively compared to control embryos, demonstrating a stark reduction in the binding of both factors in their respective mutants (Fig. 5B,C, S5A,B, Table S4). Performing spike-in CUT&Tag for Chip in *zelda^-/-^* embryos revealed almost a complete absence of Chip binding in *zelda* mutant embryos (89.5% signal reduction), which is even higher than after iLEXY blue light depletion (Fig 5C, S5C, Table S4). In contrast, Zelda occupancy is largely unperturbed after Chip depletion (Fig 5C, S5D, Table S4). These results indicate that Chip recruitment requires Zelda binding at 2-4h of embryogenesis, however, Zelda binding is largely independent of Chip.

This suggests that Chip may act as a cofactor of Zelda. To assess this, we compared the gene expression changes after Chip depletion to those in *zelda^-/-^* germline clone embryos at the same stage (2-3h) of embryogenesis (using a published *zelda* dataset^40^). To directly compare expression changes, we reanalyzed the *zelda* expression data^40^, which revealed 2,519 differential genes using the same cut offs (FDR < 0.01 and |log2FC| > 1) as we used for Chip (Table S3). Considering only the genes detected as expressed in both experiments (6,364 genes), revealed highly concordant changes. Despite causing a smaller magnitude of effect on transcription, Chip deregulated genes are a subset of those deregulated in *zelda^-/-^* mutant embryos (87% overlap. Fig. S5F) and a high correlation (*r* = 0.66) when comparing the direction of fold changes (Fig. 5D and S5G). The shared occupancy as well as the large overlap of deregulated genes provides strong support that Chip is a cofactor of Zelda at these early stages of embryogenesis.

Although Chip is not required for Zelda binding at the vast majority of its sites (Fig. 5B), there are 388 regions (out of 9286 co-bound regions) where Zelda binding is significantly reduced (|log2 fold change| > 0.5 and FDR < 0.05) in the absence of Chip, suggesting that Chip might help to stabilize Zelda binding at a small subset of their common targets. As Zelda is an essential regulator of ZGA in the early embryo^37–39^, this could be another mechanism by which Chip influences this process. To explore this, we looked at Zelda binding at the promoters of the 724 genes that are deregulated after Chip depletion at 2-4h. This revealed that 18.5% (56) of downregulated genes also have significantly depleted Zelda binding at their promoter, compared to only 2.7% (11) of upregulated genes (Fig. S5H). Interestingly, of the 107 genes identified as pre-mid blastula transition expressed genes^41^, which coincides with the major wave of ZGA, 38 have significantly decreased Zelda promoter binding after Chip depletion, indicating that for a subset of the early wave of ZGA, Chip is required to stabilise the occupancy of Zelda (Fig S5I).

Taken together, these results indicate that Chip and Zelda co-operate in regulating gene expression during early stages of embryogenesis. Zelda is required for Chip recruitment to both enhancers and promoters. Both factors therefore co-bind to the same targets, and have overlapping transcriptional responses after the removal of either factor, indicating that Chip is required to mediate part of Zelda’s transcriptional response. At the majority of sites, Chip most likely does this by interacting with cofactors or the basal transcriptional machinery, which we explore below, while at a small subset of sites it may also help to stabilise Zelda binding, especially for genes involved in the early wave of ZGA.

### Chip depletion does not affect chromatin conformation at early stages of embryogenesis

The mammalian homolog of *Chip*, *Ldb1*, can regulate gene expression in mice and humans by forming chromatin loops between an enhancer and promoter^13,15,16,27,42–44^. To investigate Chip’s role in genome architecture and chromatin loops in *Drosophila*, we performed Hi-C and Capture-C in dark and blue light depleted *chip::iLEXYs* embryos, focusing on the critical 2-4h time window where Chip is essential for embryogenesis. Surprisingly, this revealed no major changes in Topologically Associating Domains (TADs) or contact frequency. The *hairy* gene, for example, has altered expression losing some strips upon blue light depletion (Fig. 3E). However, there is no difference in contact frequency after Chip depletion, observed by either Hi-C (Fig. 6A, top) or Capture-C, using the *hairy* promoter as bait (Fig. 6A, bottom). Focusing on all enhancers and promoters bound by Chip, revealed no difference in loop intensity between dark versus blue light depletion, as observed from loop pileups (Fig. 6B) and their quantification (Fig. 6C) using the Hi-C data. Further stratification of different subsets of loops, including high-confidence manually annotated loops^45^, loops where a loop anchor is bound by Chip at a promoter, or bound by Chip at an enhancer, or overlapping a promoter of a differentially expressed gene, also revealed no detectable change in loop interactions (not shown). Moreover, insulation scores at TAD boundaries were not affected after Chip depletion (Fig. 6D).

**Figure 6:**
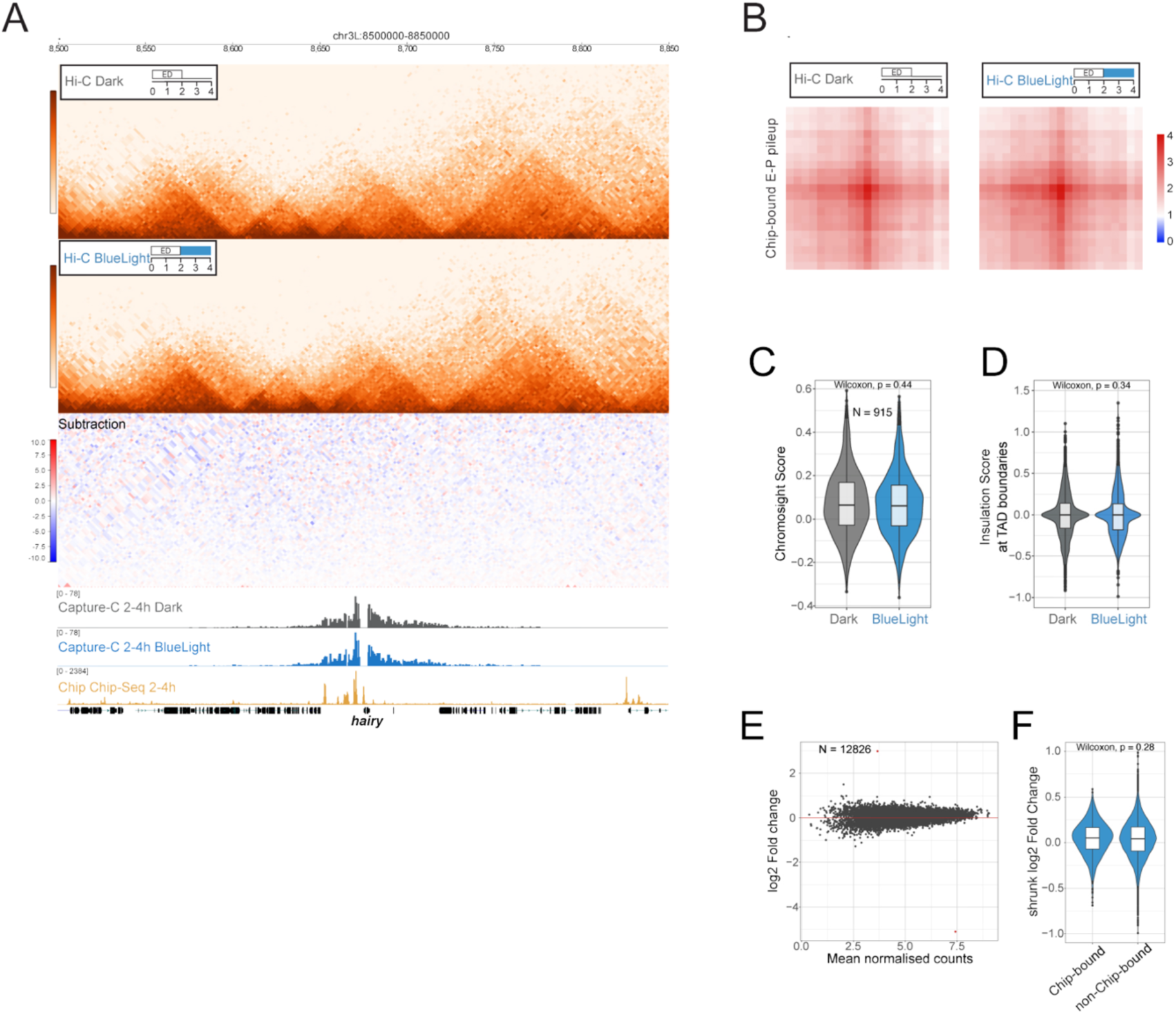
Chip depletion does not affect chromatin architecture or loops in 2-4h embryos. (**A**) *Top*: Hi-C contact matrices in 2-4h *chip::iLEXYs* embryos in dark and after 2h blue light exposure, showing a ∼350kb region centered on the deregulated *hairy* gene. The subtraction matrix, underneath, shows little differences after depletion. *Bottom*: Capture-C tracks with the viewpoint (bait) at the *hairy* promoter in embryos in the dark (gray) and blue light depletion (blue). Chip occupancy (CUT&Tag signal, yellow) in 2-4h embryos is shown underneath. (**B**) Pile-up plot of Hi-C signal at enhancer-promoter (E-P) loops defined as interactions between a TSS-Proximal Chip peak and a TSS-Distal Chip peak, in dark and blue light exposed embryos. (**C**) Violin plot of the distribution of the chromosight scores for enhancer-promoter Chip-bound loops (as in (B)) in dark and after blue light exposure. N = number of loops. (D) Violin plot of the distribution of insulation scores at TAD boundaries in dark and blue light exposed embryos. Significance calculated using the Wilcoxon rank-sum test. (**E**) MA plot of Capture-C signal in 2-4h *chip::iLEXYs* embryos in dark versus blue light exposure. The capture C baits targeted a collection of Chip-bound, Zelda-bound and control proximial and distal regions. N = number of interactions. (**F**) Violin plots of the distribution of changes in interaction frequency (log2 fold change) detected by Capture-C separated by regions bound or not bound by Chip. In panels C,D and F, the boxplots inside the violin plots display the median, 25th, and 75th percentile (hinges), whiskers and outliers (data points beyond whiskers).

To ensure that the resolution of the Hi-C data is not masking more subtle changes in chromatin architecture, we performed Capture-C in *chip::iLEXYs* 2-4h embryos after exposure to blue light for two hours, in biological replicates which were sequenced to high depth obtaining ∼5-8 million valid unique reads per replicate. We selected 501 loci for the Capture-C, which includes 97 promoters and 112 enhancers bound by both Chip and Zelda, 90 enhancers bound exclusively by Chip, 107 enhancers bound exclusively by Zelda and 95 control regions (Table S5). This revealed only 2 significant changes in contact frequency after Chip depletion out of 12,826 interactions tested (Fig. 6E). Moreover, there is no significant difference in contact frequencies between Chip-bound versus unbound regions after Chip depletion (Fig. 6F). Although we cannot exclude that the lack of detected changes in E-P loops is not due to heterogeneity already forming within the 2-4hr embryos, which could obscure cell-type specific loops, we and others have shown that many E-P loops are already formed and show little differences between cell types during these early stages^46–49^, and even at 6-8h^47^.

Taken together, those results suggest that Chip may not be required for the formation of enhancer-promoter loops at this stage of embryogenesis, even though it is essential for viability. This indicates that Chip is regulating gene expression and embryonic progression through a mechanism other than chromatin conformation at early stages of embryogenesis.

### Chip acts as a Zelda cofactor to recruit CBP and establish H3K27ac deposition at enhancers and promoters during early embryogenesis

Zelda functions as a pioneer factor during ZGA, increasing chromatin accessibility so that other TFs can bind their enhancers^50,51^, and is essential to regulate the early wave of zygotic transcription^37–40,52,53^. To determine whether Chip acts as a cofactor of Zelda in its function to establish accessibility, we performed spike-in ATAC-seq in Chip nuclear depleted embryos as well as in *zelda^-/-^* germline-clone embryos at 2-4h. Chromatin accessibility is dramatically decreased in *Zelda^-/-^* 2-4h embryos, as expected (Fig. 7A, Table S6). In contrast, there is no major change in accessibility after Chip depletion (Fig. 7B, Table S6). We confirm that in *zelda^-/-^* embryos, the sites with the largest decrease in accessibility are regions where Zelda occupancy is also significantly decreased, both at promoters (TSS proximal) and distal regions (Fig. 7C). Unexpectedly, accessibility was even slightly increased in regions where Chip occupancy is significantly decreased: 4.7% and 7.7% median signal increase at promoters and distal regions, respectively, in Chip-bound versus non bound regions (Fig. 7D). These results indicated that Chip is not required for Zelda’s function to open chromatin.

**Figure 7:**
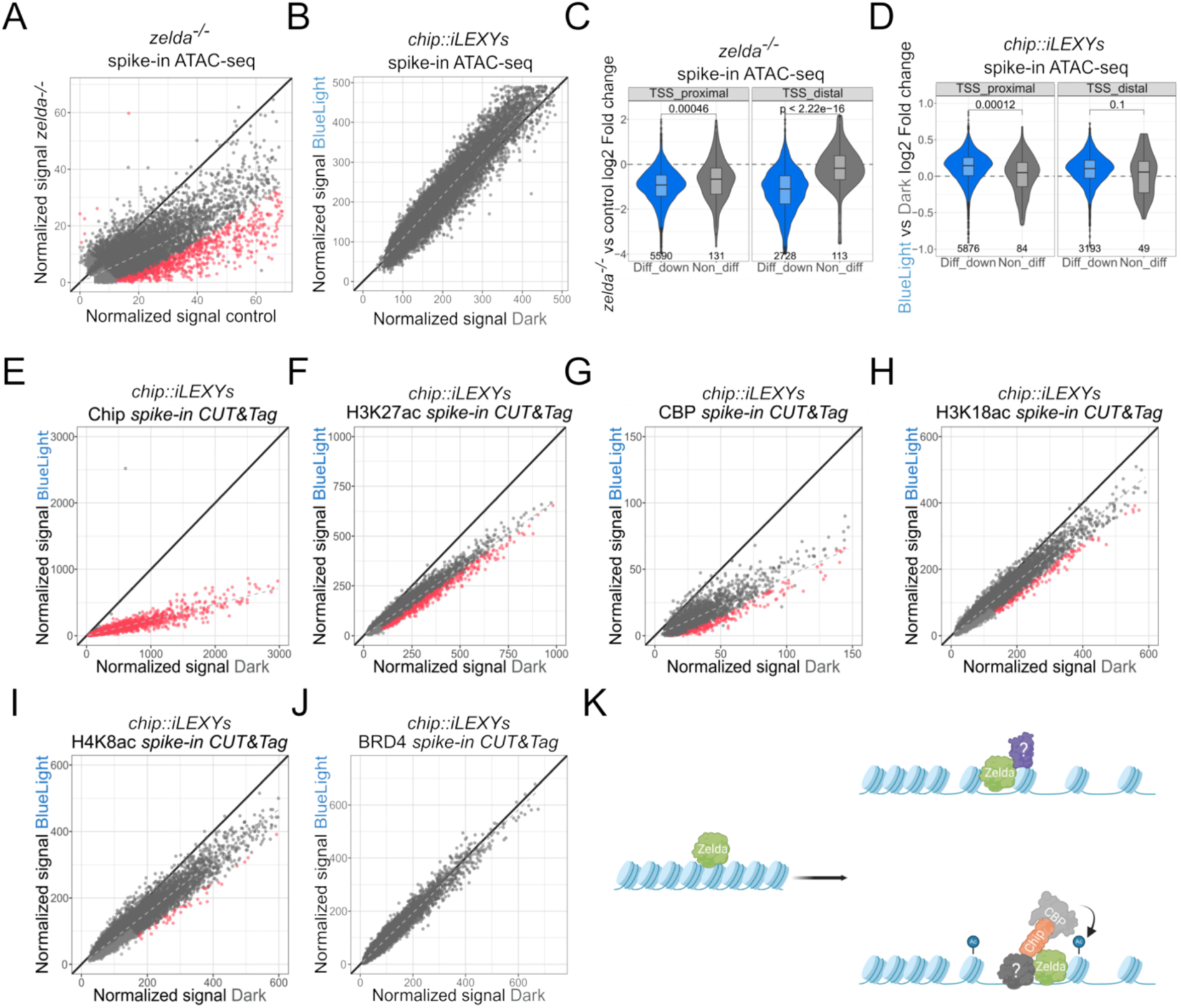
Chip is a new cofactor of Zelda in the establishment of H3K27ac through the recruitment of the coactivator CBP. (**A-B**) Scatterplots of spike-in normalized ATAC-seq signal in control versus *zelda^-/-^* germline clone embryos (**A**) and in *chip::iLEXYs* dark versus blue light exposed embryos (**B**) at 2-4h. Peaks that are significantly different (|log2 fold change| > 0.5 and FDR < 0.05) are indicated in red. (**C-D**) Violin plots of the distribution of spike-in normalized ATAC-seq peak signal (log2 fold change) in *zelda^-/-^* versus control embryos, in regions overlapping differential, non-differential and non-bound Zelda peaks (**C**) and in *chip::iLEXYs* embryos in dark versus blue light depleted embryos, in regions overlapping differential, non-differential and non-bound Chip peaks (**D**). Log2 Fold changes were compared using the Wilcoxon rank-sum test. (**E-J**) Scatterplots of spike-in normalized CUT&Tag signal for Chip (**E**), H3K27ac (**F**), CBP (**G**), H3K18ac (**H**), H4K8ac (**I**) and BRD4 (**J**) in 2-4h *chip::iLEXYs* embryos in dark versus blue light exposure. Differential peaks (|log2 fold change| > 0.5 and FDR < 0.05) are indicated in red. (**K**) Model Zelda (green) binding to closed chromatin, followed by chromatin opening through the recruitment of an unknown factor (question mark). Zelda recruits Chip, either directly or indirectly, and Chip recruits CBP to establish acetylation of the lysine 27 of histone 3. Schematic was created with Biorender.com.

Since Chip is not required for Zelda’s ability to increase accessibility (Fig. 7A-D), we next assessed if Chip is essential for a transactivation function at enhancers or promoters by performing spike-in CUT&Tag for a number of factors in *chip::iLEXYs* 2-4h embryos that developed in the dark or in 2 hours blue light exposure. Given that Zelda binding is associated with H3K27ac^54^, we first determined if there are changes in H3K27ac after Chip depletion. This revealed 722 peaks with a significant reduction (|Fold Change| > 0.5 and p-value < 0.05, Methods) upon Chip depletion, suggesting that Chip is downstream of, and likely a cofactor of, Zelda in establishing H3K27ac (Fig. 7E, F, Table S4). Regions with depleted Chip and Zelda peaks have the most significant reduction in H3K27ac signal at both TSS proximal and distal regions (Fig. S6A, B), suggesting a direct relationship between the two. We next assessed if Chip could act as a mediator between Zelda and CBP (the single *Drosophila* homolog of the mammalian CBP and p300^55,56^), which acetylate lysine 27 of H3^57,58^. CBP was recently shown to be required for ZGA in *Drosophila*^59^ and similar to Zelda, is required to establish RNA Pol II clusters in the early *Drosophila* embryo^60^. Depletion of Chip leads to a dramatic reduction in CBP occupancy (Fig. 7G, Table S4), accounting for a 52.6% global decrease in CBP signal, indicating that Chip is required for CBP recruitment. As CBP can acetylate other histone lysine residues, we also examined H3K18ac and H4K8ac signal after Chip depletion, which revealed a less profound effect, with 172 and 46 peaks significantly decreased, respectively (Fig. 7H, I, Table S4). We also examined Brd4 (also called *fs(1)h*) occupancy, which binds to acetylated histones through its bromodomains and regulates the transcription of a subset of genes^61,62^. Despite the significant decrease in H3K27ac signal, we did not observe significant changes in Brd4 binding upon Chip depletion (Fig. 7J, Table S4), which was also observed in another study^63^. This was surprising as Zelda and CBP were reported to nucleate Brd4 into clusters or foci observed by imaging^60^, and suggests that Zelda can influence Brd4 nuclear localization by a Chip (Ldb1) independent mechanism.

Collectively, these data indicate that Chip (Ldb1) acts as a functional bridge between Zelda and CBP to establish H3K27ac at both promoters and (putative) enhancers in early embryos during ZGA (Fig. 7K).

## DISCUSSION

Since its discovery in *Drosophila* almost 30 years ago, Chip (Ldb1) function has been linked to many aspects of gene regulation. This includes enhancer-promoter communication in *cis*, in both *Drosophila*, mouse and human loci, interchromosomal interactions at the mouse olfactory loci, and activity-related function at enhancers and promoters. Despite these many interesting roles, there has been surprisingly little genetic dissection of the function of Chip (Ldb1), except at a handful of model loci; *cut*^11,12^ and *achaete/scute* (ac/sc)^24,64^ genes in *Drosophila*, *α-* and *β-globin* loci in mice and human^13,14,19,27–31,65,66^, and the olfactory cluster in mouse^17,26^. No genome-wide assessment had been performed to date that could generalized these functions, with the exception of a recent preprint using auxin-degron in a mouse erythroid cell line^67^. There has been no investigation of the requirement of Chip (Ldb1) in the context of embryogenesis, even in *Drosophila* where the gene was first discovered.

This is mainly due to the technical challenges of dissecting the role of cofactors, which by design are very pleiotropic. Cofactors tend to be recruited broadly across the genome, and are active in almost all cell types, being typically ubiquitously expressed and maternally deposited. To study their function, the maternally deposited protein must be removed, which is typically done in *Drosophila* by making germline clones (as we did here for *zelda*), or using knockdown in the oocyte by RNAi. For many cofactors, this is not possible, as the factor is required for the process of oogenesis itself – BRD4 removal from the germline, for example, leads to female sterility giving rise to the gene’s name *fs(l)h*^68–70^. When mutant embryos can be obtained, our understanding of the function of pleiotropic factors in embryogenesis is usually limited to the very early stages, as development becomes blocked at the first time-point of requirement. To circumvent these issues, we applied the iLEXY system to deplete Chip (Lbd1) from the nucleus (including the maternally deposited protein) at precise time-windows of embryogenesis, assessing its role on transcriptional regulation, and its necessity for viability, for the first time at different stages of embryogenesis. Importantly, iLEXY activation very efficiently depletes Chip from the nucleus, removes it from chromatin and mimics the phenotype of germ-line *loss-of-function* mutant embryos. This revealed that Chip (Ldb1) has distinct molecular functions at different stages of embryogenesis.

The speed and reversibility of the iLEXY system allowed us to narrow down the critical window for viability to just one hour in embryogenesis, despite Chip being expressed through all stages. These results highlight a new level at which one can dissect the function of a nuclear factor at unprecedent temporal resolution in a developmental setting, uncovering stage-specific molecular and organismal phenotypes.

### The function of Chip during early embryogenesis appears independent of a role in chromatin looping

Classic genetic analysis in *Drosophila* showed that Chip (Ldb1) is involved in E-P communication at the *cut* locus during wing margin development^11,12^ and at the *ac*-*sc* complex during proneural patterning^24,64^. In both cases, interactions between a Chip dimer and the LIM domain-containing TF, Apterous, was thought to form a tetramer that could explain the E-P interactions^71,72^. However, both studies lacked direct evidence showing that Chip establishes a chromatin loop between the enhancer and promoter, as no such technology existed at the time.

In mammals, Ldb1 mediates looping at the *β-globin*, *Myb* and *Epb4.2* loci in erythroid cells^13,27,42^ in addition to other loci in hepatocytes and myogenic lineages^43,44^. Artificial tethering of Ldb1 to the promoter of *β-globin*, in the absence of the activator GATA-1, can form a loop between the LCR and the promoter through interactions with the endogenous Ldb1 complexes and restore expression^15^. Ldb1 is also required to orchestrate enhancer interactions in mammalian olfactory receptor choice, in this case bridging enhancers in both *cis* and in *trans*. Interestingly, a similar regulatory module (the TF Lhx2 recruiting Ldb1) is used in a different context in *Drosophila* (Apterous-Chip, the *Drosophila* homologs of Lhx2-Ldb1)^17^.

Given all of this, we were very surprised to observe no changes in chromatin interactions after Chip depletion. Even using high-resolution Capture-C focusing on enhancers and promoters with changes in Chip binding and/or expression after Chip depletion, and performing the experiments at the stages where Chip (Ldb1) is essential for embryogenesis (2-4hr), we did not observe any obvious changes in E-P looping. We cannot exclude that we still don’t have high enough resolution. In the preprint by Aboreden et al^67^, they used Micro-C to detect loops that change after auxin-mediated Ldb1 degradation. This revealed only a relatively small fraction of loops that decreased in strength: ∼909 out of 3970 Ldb1-bound loops. The authors state that using a higher resolution method (Region Capture Micro-C) could uncover even more Ldb1-dependent loops. However as the method will also uncover many more unchanged loops, this will still represent a small fraction of Ldb1-bound loops. This suggests that Ldb1 may regulate E-P looping at a small subset of specific genes, being brought there by the appropriate TF for that cell type and stage. Extrapolating to *Drosophila*, with a genome ∼25-fold smaller, it is not clear how many loops we could expect to change, assuming the gene have the same function.

We selected early embryogenesis (2-4h) for the chromatin conformation capture experiments as this is the window when Chip is essential for viability. However, we and others, have shown that at these early stages, enhancers and promoters generally act over quite short distances, and are present in pre-formed topologies before the gene is activated and even before ZGA^46–48^. This is in contrast to later stages of embryogenesis, where there are new, often more distal enhancer and promoter interactions^47,73^. The imaginal discs, where the role of Chip was first described, are post-embryonic structures, developing during larval stages and may have more distal regulation. At least this holds true for the *cut* wing-margin enhancer, which is ∼80kb away from its target gene^11,12^. It will be very interesting in future studies to explore the role of Chip (Ldb1) in chromatin looping at other stages of *Drosophila* embryogenesis or in adult tissues, especially in imaginal discs where its role was first discovered.

### Chip is a functional bridge between Zelda and CBP during ZGA

Our genetic dissection of Chip function, shows that it is essential for viability in a narrow window that overlaps the ZGA. We show that Zelda recruits Chip to chromatin, and subsequently Chip recruits CBP to these Zelda bound sites. The depletion of Chip leads to extensive changes in H3K27ac at both enhancers and promoters, and changes in gene expression that mirror *zelda* mutant embryos. Initially Chip, and its vertebrate homolog Ldb1, were thought to be recruited to chromatin mainly through LIM domain factors, hence the name <u>L</u>IM <u>d</u>omain <u>b</u>inding protein 1 (Ldb1)^26,74–77^. Chip can homodimerize, and binds to TFs that either bind to DNA as homodimers (i.e. LIM domain TFs, e.g. Apterous dimer) or heterodimers (e.g. Ac/Da), forming a functional tetramer. *Chip loss-of-function* germline clones leads to a segmentation defect. However, there are no known LIM domain TFs that can phenocopy those defects, suggesting that at these early stages, Chip is recruited by other factors^12^. Chip interacts physically with - and in some cases is recruited to chromatin by – other TFs, including those belonging to the homeodomain-, GATA-, bHLH-family of factors^18,24,26,28,74,77–81^. Chip also physically interacts with the architectural proteins CTCF^65^ and Su(Hw)^78^, highlighting many ways in which it can be recruited to chromatin. However, during this 2-4hr window, Zelda is the main recruiter of Chip, as there is almost a complete loss of Chip binding in *zelda* germline clone embryos (Fig. 5C).

Chip is also a characterized component of the Wnt (*wg* in *Drosophila*) enhanceosome, and through interactions with different TFs and cofactors, Chip can confer context specificity^18,81,82^. In the Wnt OFF state, the corepressor Groucho is bound to enhancers, while in the Wnt ON state this is replaced by CBP, which induces histone acetylation^18,81^. Chip is therefore present in both repressive and activating complexes, which could explain the down and up -regulated changes in gene expression that we observe upon Chip depletion, and emphasizes Chip’s versatility as a cofactor or scaffolding protein. CBP also acetylates lysines in other nuclear proteins (not only histones), and this acetylation can regulate their function - acetylation of TCF, for example, disrupts its interaction with Armadillo (a TF downstream of Wg signaling) and thereby negatively regulates Wnt signaling^83–85^. Catalytic mutants of CBP appear to rescue viability in embryos that are mutant for CBP and can activate zygotic gene expression^59^. This suggests that the recruitment of CBP by Chip (Ldb1) in early embryos will likely have additional functions (independent of H3K27ac) in complexes that are unknown to date.

### Two ‘molecularly’ independent functions of Zelda during ZGA

Several processes must be tightly regulated and coordinated for ZGA to occur correctly to drive this crucial stage in early embryogenesis: an increase in chromatin accessibility for thousands of regions, the deployment of certain histone modifications to help activation, activation of the RNA polymerase II activity at various steps (release from pausing, elongation), as well as the decay of the maternally loaded RNA transcripts^86^. Zelda appears to orchestrate many of these processes and has multiple functions at distinct regions^40^, suggesting that it may associate with different sets of cofactors. The pioneering role of Zelda to open chromatin may require the recruitment of a chromatin remodeling complex, for example, although this has not been identified to date^87^. In some genomic regions, Zelda acts through a functional association with other pioneer factors such as CLAMP^88^ or GAF^89^. However, the pioneering function of Zelda (and the recruitment of a cofactor) does not require Chip (Ldb1), as Zelda can still open chromatin after Chip depletion. Interestingly, the quantitative CUT&Tag even indicates the opposite - a small, but significant increase in accessibility at Chip bound regions after Chip depletion. As almost all of these regions are cobound by Zelda, we speculate that there may be two distinct Zelda complexes that in certain regions of the genome, might be weakly competing. As a result, in the absence of Chip, Zelda recruits another cofactor and accessibility increases in those regions.

Our data indicates that the primary role of Chip (Ldb1) as this stage of embryogenesis is to help Zelda, not in its pioneering role, but rather to mediate (at least some of) its transcriptional activating role. It was recently suggested that CBP might be a cofactor of Zelda since both are required for RNA Pol II clustering into hubs during ZGA and CBP is genetically downstream of Zelda in this activity^60^. Our data uncovers Chip as the missing cofactor connecting the two, regulating CBP recruitment at a fraction of Zelda-bound sites. Whether this takes place in a single complex with direct protein-protein interactions or through the recruitment of those factors via different TFs to the Zelda-opened regions remains unknown (Fig. 7K).

How the specificity between the different functions of Zelda and the different regions in the genome is achieved remains unclear. While almost all of Chip’s function at 2-4h appears to be mediated through Zelda, as seen from the almost complete overlap in binding, only part of Zelda’s function (although a majority) are mediated through Chip. From their co-occupancy, 71% (9,286/13,097) of the significant Zelda peaks are co-bound by Chip, with many of the remainder being sub-threshold levels of Chip (Fig. S5E). However, there are clearly additional cofactors of Zelda yet to be discovered.

### Limitations of the study

Our ChIP-seq and RNA-seq data indicate that Chip has different functions at different time-windows of embryogenesis. Although there is no obvious difference in the level of Chip depletion by immunostaining at 6-8h and 10-12h compared to 2-4h (Fig. S3), we note that we did not assess the depletion of Chip from chromatin at these later stages. There is therefore a possibility that Chip is not depleted as efficiently from chromatin in later embryos, especially at 10-12h, where we observe only minimal differences in gene expression. However, we think this is unlikely based on three observations: First, Chip has very weak binding at 10-12h in control embryos both at enhancers and at promoters (Fig. 1C), suggesting that it may not have a functional role at this stage. Second, we do observe many transcriptional changes after Chip deletion at 6-8h, indicating that it is depleted (Fig. 3A). Third, we have observed very similar chromatin depletion for other factors at later stages (6-8h and 10-12h) compared to earlier stages (2-4h), indicating that the increase complexity of the embryo itself is not a major factor.

We provide extensive data that demonstrates that Chip has an essential role for Zelda’s function and for early embryogenesis: the requirement of Zelda for Chip recruitment, their very extensive co-binding and their concordant changes in gene expression in both mutants, provides very strong evidence that Chip is a new cofactor of Zelda. The essential critical window of Chip, overlapping the ZGA, for embryogenesis further strengthens these findings. However, we still don’t know if Zelda, Chip and CBP belong to a single complex, with physical protein interactions between each other or alternatively, if Zelda first opens chromatin and then some other TF binds, recruits Chip which in turn recruits CBP, where three factors form a complex that doesn’t include Zelda (Fig. 7K). Our data is compatible with both scenarios. A homeodomain motif (Dve, Oc) was the second most enriched motif at Chip regions specifically bound a 2-4h (Fig. 1D). We tried to co-IP Chip and Zelda, but got variable results, so this is still unclear. Future biochemical experiments combined with precise functional perturbations, made possible by optogenetics, are going to be invaluable to distinguish between these two possibilities.

## AUTHOR CONTRIBUTIONS

C.C.G., Y.K. and E.E.M.F. conceptualized and planned the study. Y.K. performed the CRISPR experiment and embryonic viability assays; C.C.G. and Y.K. performed embryo collections; C.C.G. and R.M.-F. performed in-situ hybridization; C.C.G., Y.K. and R.R.V. performed genomics experiments; M.F. performed bioinformatic data analyses. C.C.G. and E.E.M.F. wrote the manuscript with input from all authors. E.E.M.F. supervised the project and provided funding.

## ACKNOWLEDGMENTS

The authors thank all Furlong lab members for very helpful comments. We are grateful to Dale Dorsett, Christine Rushlow and Mattias Mannervik for sharing reagents. This work was technically supported by EMBL’s Genomics Core, Advanced Light Microscopy and Flow Cytometry Core facilities, and by the external resources provided by Bloomington Drosophila Stock center and Flybase. We are very grateful for financial support from EMBO (post-doctoral fellowships awarded to C.C.G. (297-2021) and Y.K. (1098-2018)), and Deutsche Forschungsgemeinschaft (DFG SPP 2202) and European Research Council (ERC advanced grant) agreement 787611 (DeCRyPT) grants awarded to E.E.M.F.

## Supplemental Information

**Figures S1-S6**

**Tables S1-S6**

## METHODS

### EXPERIMENTAL MODEL AND STUDY PARTICIPANT DETAILS

*Drosophila melanogaster* (*D*. *melanogaster*) and *Drosophila virilis* (*D*. *virilis*) were maintained in a standard cornmeal-agar medium and grown at 25°C. The *zld^294^*allele (gift from Christine Rushlow) was used to create *zelda* germline clones as previously described^37^. As a control, stage matched embryos were collected from y[1]w[1] (*yw*) line (Bloomington stock 1495). The following line was used generate the CRISPR-Cas9 mediated knock-in of the iLEXYs tag to the *chip* locus: w[1118]; Pbac{y[+mDint2] = vas-Cas9}VK00027 (*vasa-Cas9*) (Bloomington stock 51324).

### METHOD DETAILS

#### CRISPR tagging *chip* with iLEXYs

To tag the endogenous *chip* locus with the iLEXYs cassette^35^ the scarless CRISPR-Cas9 gene editing technology (https://flycrispr.org/scarless-gene-editing/) was used as follows: gRNAs targeting sites close to the *chip* stop codon were designed using the flyCRISPR Target Finder (tools.flycrispr.molbio.wisc.edu/targetFinder/)^90^. The annealed oligos Chip-gRNA_for: CTTCGAAGTTCTATTGCGATACAA and Chip-gRNA_rev: TTGTATCGCAATAGAACTTCCAAA corresponding to gRNA sequence were inserted into the BbsI site of the pU6-BbsI-gRNA vector^90^. A fusion product of the *chip* left homology (LH) arm (1503 bp region directly upstream of the STOP codon, amplified from the targeted fly line), the iLEXYs fragment, and the genomic region between the gRNA cleavage site and the TTAA sequence were inserted into the AarI site of the pHD-DsRed-Scarless donor vector^90^. The *chip* right homology (RH) arm (spanning the TTAA and its 1453 bp downstream sequence) was amplified from the targeted fly line and integrated into the SapI site the same plasmid.

CRISPR donor and gRNA plasmids were combined at a 2:1 molar ratio (Donor:gRNA) and injected into *vasa-Cas9* embryos by EMBL’s *Drosophila* injection service. Positive transformants were identified by DsRed expression in the eyes of adult F1 flies and used to establish stable stocks. The precise integration of the tag and the integrity of the *chip* locus were confirmed by PCR amplification and subsequent Sanger sequencing of the CRISPR allele, from outside of the left homology arm to outside of the right homology arm^90^.

#### Blue light induced nuclear depletion

Blue light illumination of staged *D. melanogaster* embryos was performed using a custom-made, programmable LED-based blue light illumination box, as previously described^35^. Unless otherwise specified, 2-second blue light pulses at 70% intensity were alternated with 1-second breaks.

#### Embryo viability assays

Embryo viability was assessed by determining the hatching rate of embryos into first instar wandering larvae. After three 1-hour pre-lays, embryos were collected at 25°C on apple juice agar plates and, if necessary, aged to the desired stage. Yeast paste was removed from the apple plates, which were then incubated in the LED box under blue light illumination or kept in the dark at 25°C. The number of hatched and unhatched eggs was counted to determine the embryo viability rate. Importantly, all procedures were conducted under dark (safelight) conditions unless otherwise specified, and embryos were only exposed to blue light at the designated times.

#### Immunostaining and in-situ hybridisation

Immunostainings and RNA in-situ hybridization were performed as previously described^35^. The rabbit anti-Chip and the mouse anti-Lamin (DSHB #ADL101) antibodies were used at 1:100. the Plan-Apochromat 20x/0.8 M27 air (FWD=0.55mm) and 63x/1.4 Oil DIC M27 objectives.

The in-situ probes were amplified from a previously cloned plasmid for *eve*, and from the corresponding EST clones (from DGRC) for *hb* (#LD34229), *tll* (#IP01133) and *hairy* (#RE40955) using Digoxigenin- or Fluorescein-modified nucleotides. Probes were detected using the TSA Plus Fluorescence kit (PerkinElmer #NEL760001KT) after incubating with either the anti-Digoxigenin-Peroxidase (Roche #11633716001) or the anti-Fluorescein-Peroxidase (Roche #11426346910) diluted to 1:2000. Embryos were mounted in ProLong^TM^ Gold with DAPI (ThermoFisher Scientific #P36931). The immunostains were imaged on a Zeiss LSM 780 using the Plan-Apochromat 20x/0.8 M27 air (FWD=0.55mm) and 63x/1.4 Oil DIC M27 objectives. In-situs were imaged on a Zeiss LSM 880 AiryFast using the Plan-Apochromat 20x/0.8 M27 air (FWD=0.55mm) objective.

#### Quantification of Chip nuclear to cytoplasmic ratio after blue light depletion

The nuclear and cytoplasmic fraction of Chip in *chip:iLEXYs* line upon blue light illumination were quantified from immunostaining of Chip. Anti-Lamin staining was used to determine the position of nuclei. The area and the RawIntDen of manually drawn regions of interest (ROI) in the cytoplasm and the nucleus of each cell were measured using Fiji and these values were used to calculate the nuclear/cytoplasmic fluorescence ratios.

#### Fixation for ChIP-seq, CUT&Tag, Hi-C and Capture-C

Staged *D. melanogaster* and *D. virilis* embryos were collected and fixed as previously described^91^. In brief, embryos were collected in staged two-hour or one-hour windows after three one-hour pre-lays to clear the females and synchronize the collections. The collections were aged at 25°C to the corresponding time window (2-3h, 2-4 h, 6-7h, 6-8 h, 10-12 h), and exposed to blue light for the indicated durations (as stated in the results). Embryos were fixed in 1.8% formaldehyde for 15 minutes, with the exception of ChIP-seq and CUT&Tag using the rabbit anti-Chip antibody and guinea pig anti-CBP antibody, where 3% formaldehyde for 30 min was used. The formaldehyde was quenched with 125 mM Glycine in PBT (1x PBS + 0.1% Triton-X100). After two washes in PBT, the embryos were dried on a Nitex membrane, snap-frozen in liquid nitrogen and stored at −80°C until further use.

#### Nuclei extraction for ChIP-seq, CUT&Tag, Hi-C, Capture-C and ATAC-seq

Nuclear isolation was performed as described in ^92^. For ChIP-seq, nuclei were sonicated (12 cycles 30’’ ON/ 30’’ OFF) using Bioruptor Pico (Diagenode) to generate 250-500 bp chromatin fragments. The chromatin was centrifuged at 16.000 g and the supernatant containing the chromatin was aliquoted into fresh tubes and stored at −80°C until use. We used 25 μg of chromatin as input and followed the protocol as in^93^. For CUT&Tag, Hi-C, Capture-C ATAC-seq, nuclei were quantified after DAPI staining using a FACSymphony^TM^ A3 cell analyzer together with CountBright Absolute Counting Beads (Thermo scientific C36950) at the EMBL Flow Cytometry Facility. For CUT&Tag and ATAC-seq aliquots of 2 million nuclei were snap-frozen in 25 μl nuclear freezing buffer (50 mM Tris pH 8 / 25% glycerol / 5 mM MgAc2 / 0.1mM EDTA / 5 mM DTT). For Capture-C, 5 million nuclei were snap-frozen without any liquid.

#### Spike-in CUT&Tag

Nuclei were thawed on ice and resuspended in PBT. DAPI was added and single nuclei were sorted using the BD FACSAria^TM^ Fusion. 50 thousand *D. melanogaster* together with 50 thousand *D. virilis* nuclei were sorted into ice-cold dig-wash buffer (20 mM HEPES pH 7.5, 150 mM NaCl, 0.5 mM Spermidine, 0.01% Digitonin, 1x Roche cOmplete Protease inhibitors) and kept on ice. The CUT&Tag was performed as described previously using prewashed Concanavalin A beads (Polysciences Europe GmbH, #86057-3)^93^. In brief, for tagmentation, nuclei were resuspended in 200 μl of CUT&Tag Tagmentation buffer (36.3 mM Tris-Acetate pH 7.8, 72.6 mM K-Acetate, 11 mM Mg-Acetate, 17.6% Dimethyl-Formammide) and incubated at 37°C for 1h. Tagmentation was stopped and samples de-crosslinked (25 mM Tris pH 8, 50 mM EDTA, 0.1% SDS, 50 mM NaCl, 0.5 mg/ml Proteinase K (Qiagen 19131)). The next day, DNA was purified by column purification (Zymo, DNA clean and Concentator-5 Kit, #D4014). The samples were then RNase treated and PCR for library preparation was performed using P5 and P7 primers using the NEBNext® High-Fidelity 2x PCR Mater Mix (M0541L). The PCR products were purified with 1.3x Agencourt AMPure XP beads (Beckman Coulter, #A63881), quantified with Qubit, ran on the Bioanalyzer using hs DNA reagents and chips and used for sequencing.

The primary antibodies used for CUT&Tag and ChIP were as follows: Rabbit anti-Chip antibody (serum was a gift from Dale Dorsett, which we affinity-purified), Rabbit anti-Zelda (gift from Christine Rushlow), Rabbit anti-H3K27ac (Abcam ab4729), guinea pig anti-CBP (gift from Mattias Mannervik), Rabbit anti-H4K8ac (Abcam ab15823), Rabbit anti-H3K18ac (Abcam ab1191). Secondary antibodies for CUT&Tag: Guinea Pig anti-Rabbit IgG (Antibodies-online, ABIN6923140), Goat anti-Guinea Pig IgG (Antibodies-online, ABIN6923140).

#### RNA-seq, Hi-C and Capture-C

RNA-seq experiments were performed in 3-4 biological replicates per genotype as described in ^93^. For Hi-C, we used the Bridge-Adaptor *in situ* Hi-C protocol as described in^93^, in biological replicates per genotype. Capture-C was performed in biological replicates per genotype, using 25 million nuclei per replicate to obtain high biological complexity per sample. We used our established Capture-C protocol^47^, with the following modifications to the hybridization step. Pooled samples (375ng each) were processed according to the Twist Target Enrichment protocol, with minor modifications. Briefly, the indexed libraries were dried using a vacuum concentrator and resuspended in a blocker solution with 5 µl SeqCapEZ from NibleGen (#06684335001, Roche) and 7 μl universal blockers. In parallel, a probe solution was prepared by adding 4 µl of the Capture-C probes (Table S5) to the Hybridization Mix, heating at 65°C for 10 min and slowly cooling down for 5 min at RT. For the hybridization reaction, the probe solution was heated to 95°C for 2 min and cooled on ice for 5 min, the library pool was heated to 95°C for 5 min, and both were equilibrated at RT for 5 min before mixing. 30 µl Hybridization Enhancer was added and the reaction incubated at 70°C for 16 h. The hybridized library was then bound to streptavidin beads, PCR-amplified for 13 cycles and purified using DNA Purification beads. After quantification with Qubit and quality control on a Bioanalyzer (displaying a single peak between 350 and 500 bp) they were used for sequencing.

#### Spike-in ATAC-seq

For ATAC-seq, nuclei were thawed on ice and resuspended in PBT. DAPI was added and single nuclei were sorted using the BD FACSAria^TM^ Fusion. 225 thousand *D. melanogaster* together with 75 thousand *D. virilis* nuclei were sorted into PBT. Nuclei were spun down and incubated in 1 ml PBT with 1x protease inhibitors and 0.4% IGEPAL CO-630 (Sigma, 542334) for 30 minutes on a nutator at 4°C. The suspension was centrifuged for 5 min at 3200 g and 4°C and the pellet resuspended in transposition solution containing 5 μl TDE1 (Illumina, 15027865) 25 μl 2x Tagmentation DNA buffer (Illumina, 15027866), 20 μL nuclease-free H_2_O and incubated 30 min at 37 °C. 50 μl of stop solution (50 mM Tris-HCl (pH 8.0), 100 mM NaCl, 0.1 % SDS, 100 mM EDTA (pH 8.0)) and 5 μL of RNAse at 1 mg/mL were added. After incubation of 10 min at 55 °C 3 μl of Proteinase K (Qiagen 19131) was added and samples incubated for 1 h at 65 °C. DNA was purified using the DNA clean and concentrator kit (Zymo, D4014) and eluted from the column in 20 μl of EB buffer. DNA concentration was quantified by Qubit hs dsDNA, followed by amplification by PCR (12 cycles) using the Nextera DNA sample preparation and indexing kits (Illumina, FC-121-1030, FC-121-1011). DNA fragments were size-selected using 0.5 – 1.8x AMPure XP beads (Beckman Coulter, A63881) and eluted in a final volume of 15 μl. The samples were measured by Qubit and fragment size distribution checked on a Bioanalyzer DNA hs chip before sequencing.

### QUANTIFICATION AND STATISTICAL ANALYSIS

#### ChIP-seq

##### Mapping, coverage and QC

Reads were trimmed with Trim_galore (v. 0.4.3.1) requiring a minimum base quality of 20 and a minimum read length of 20. Reads were then mapped with bowtie2^94^ (v. 2.3.4.2) with options “--un -X 2000 --fr --dovetail --sensitive --no-unal”. The resulting bam file was filtered with samtools^95^ (v.1.1.2) requiring a minimum mapping quality of 20, including only reads mapped in proper pairs and removing supplementary alignments. Only reads mapping to major chromosome (Chr 2, 3, 4 and X) were kept for further analyses. Duplicate reads were removed with picard^96^ (v 2.7.1.1) MarkDuplicates with options “REMOVE_DUPLICATES”. Coverage tracks were generated with bamCompare (deeptools^97^ v. 3.0.2.0) by subtracting input signal and extending reads. Signal tracks (bigwig) were obtained using bamCoverage (from deeptools) with options “--binSize 10 --effectiveGenomeSize 125464728 --extendReads”. FastQC^98^ (v. 0.69) was used to check sequencing quality. Picard CollectAlignmentSummaryMetrics (v. 2.7.1.1) was used to evaluate the mapping quality metrices across samples (mainly read duplication and unmapped reads). Insert size distribution was assessed with Picard CollectInsertSizeMetrics (v. 2.7.1.1). Correlation between samples was checked with deeptools multiBigwigSummary and plotCorrelation^99^ (v. 3.0.2.0) based on 10bp bigwig files. QC results were summarized and compared across samples with multiQC^100^ (v. 1.13).

##### Peak calling and quantification

Peaks were called using the IDR pipeline following the ChIP-seq guidelines from Encode and modENCODE^101^ based on peak summits to increase precision of peak detection. Peaks were called with macs2^102^ (v. 2.2.7.1) with options “-p 0.6 -f BAMPE -- call-summits -g 1.2e8”. The resulting summits were expanded by 80bp upstream and downstream and tested for reproducibility with IDR^103^ (v. 2.0.3) and options “--input-file-type narrowPeak -- rank signal.value --plot -i 0.05”. IDR peaks were centered and expanded by 100bp up and downstream and filtered for having a signal enrichment of > 3 times the ChIP-seq background, to ensure that peaks corresponded only to regions with strong signal over noise ratio. Peaks across time-points were merged with DiffBind^104^ (v. 3.8.4) to create a unique peak set. The merged peaks were expanded by 300bp up and downstream and overlapped the IDR peaks using bedtools intersect^105^ (v. 2.27.1) to recover the time at which the merged peaks were bound by Chip (Fig. 1B, Table S1).

##### ChIP-seq analysis

Characterization of Chip binding profile (Fig. 1A) was obtained with the R package ChipSeeker^106^ (v. 1.34.1), based on the R package org.Dm.eg.db (v. 3.16.0). Time-point specific peaks at 2-4h, 6-8h and 10-12h (Fig. 1B,D, Table S1) were obtained by testing for differential accessibility with DiffBind. Peaks in one time-point were compared with the other two time-points pairwise, requiring them to be differentially bound and with larger signal in both contrasts, using the following stringent cutoff: |log2Fold Change| > 2 and FDR < 0.05. The signal over peaks was calculated with DiffBind using the average per replicates RPKMs (Table S1). The heatmap for time-point specific peaks (Fig. 1D) was obtained by plotting z-score normalized RPKM signal of time-point specific peaks using the R package pheatmap. The ChIP-seq signal heatmaps (Fig. 1C) were obtained with deeptools computeMatrix and plotHeatmap^99^ (v. 3.5.2).

Peaks distance to TSS (Fig. S1B) was computed from the peak summit (center) to the closest TSS. The TSS annotation was obtained from Flybase^107^ (release 6.13) by including the first bp of all features annotated as “mRNA” and “gene”. Gene ontology enrichment of genes with Chip binding at their promoter (peak within 500bp from TSS) was calculated with R package topGO^108^ (v. 2.50.0) using all expressed genes at the same time-point as background (Fig. S1C-F). The enrichment was performed on Biological Process, and the results filtered for terms with a minimum of 5 and a maximum of 500 genes and FDR < 0.01. The top 15 terms were sorted by fold enrichment. Overlap between Chip TSS-Distal peaks and annotated active enhancers were performed using a collection of characterized enhancers in transgenic embryos (CAD4 enhancers^36^: intersection of enhancers from Vienna Tiles^109^, RedFly database^110^ and experimentally tested enhancers from our laboratory^92^) (Fig. S2A-D). ChIP-seq signal on Chip (Ldb1) peaks was quantified using DiffBind to obtain counts that were then library size normalized to TPMs (Transcripts Per Kilobase Million). Enhancers were overlapped with Chip peaks and in case of multiple Chip peaks overlapping the same enhancer, the one with the highest TPM value was retained.

#### Motif enrichment

Motif enrichments were obtained by testing for known motif enrichment with ame^111^ (meme suite) (Fig. 1D, Table S2). The tested regions are time-point specific Chip peaks, separated by TSS-Proximal (promoter, <500bp of any annotated TSS) and TSS-Distal (enhancer, further than >=500bp from any annotated TSS). The control regions included all Chip peaks across all time points within the same category, excluding peaks in the test regions (e.g. time-point specific enhancers against all enhancers). All motifs from CisBp^112^ (build 1.02) were used. Test and control regions were centered on summits and expanded by 50bp upstream and downstream - ame was run with the following options “--bgformat 0 --pseudocount 0.25 --method fisher”. Only motifs with an adjusted p-value < 0.05 were considered significant (Table S2).

#### RNA-seq

##### Mapping and differential expression

The read alignment and gene expression quantification was performed with the RSEM^113^ pipeline (v. 1.3.1) with options “--paired-end --strandedness reverse --star --no-bam-output --star-gzipped-read-file” and based on Flybase 6.37 gene annotation. RSEM expected counts were rounded to the closest integer and used as input for DESeq2^114^. Genes with a mean of more than 10 counts across contrasts (e.g. *chip::iLEXYs* dark [two replicates] and blue light [two replicates] samples) were considered expressed and tested for differential expression. Differentially expressed genes were defined as FDR < 0.01 and a |log2FC| > 1 (Fig. 3A-D, Table S3). The fastq files from the external *zelda^-/-^* RNA-seq dataset^40^ were downloaded from GEO and processed following the same pipeline (Fig. 5D, S5G and Table S3).

##### Gene ontology enrichment

Gene ontology enrichment of differentially expressed genes with Chip binding at their promoters (peak within 500bp from TSS) were calculated using the R package topGO^108^ (v. 2.50.0) using all expressed genes at the same time-point as background (Fig. S4A). The enrichment was performed on Biological Process, and the results filtered for terms with a minimum of 10 and a maximum of 500 of genes and an FDR < 0.05. The top 15 terms sorted by FDR are plotted.

#### Spike-in CUT&Tag and ATAC-seq

Adding spike-in nuclei from another species to both the CUT&Tag and ATAC-seq serves two purposes: 1) the exogenous reads (*D. virilis*) can be used to spike-in normalize the endogenous (*D. melanogaster*) signal and thereby allow for a quantitative comparison. 2) The exogenous signal can be used to verify that the CUT&Tag experiment worked as expected, especially in samples with extreme depletion (e.g. Chip antibody after Chip depletion) that would otherwise be comparable to a failed experiment without a proper internal control.

##### Mapping and coverage tracks

As the spike-in CUT&Tag libraries contain reads from both *D. melanogaster* and *D. virilis,* reads were mapped to a joint genome concatenating the chromosomes of both species. Less than 1% of reads mapped to both species demonstrating the large divergence between the two species genomes^115^. Only reads with unique alignments in one species were retained. Reads were trimmed with Trim_galore (v. 0.6.7) using default settings and mapped with bwa mem^116^ (v. 2.3.4.2) to a joint genome of *D. melanogaster* (dm6) and *D. virilis* (r1.07) obtained by concatenating the fasta files. The resulting bam file was sorted with samtools^95^ (v. 1.16.1), and duplicates marked with picard^96^ (v. 3.0.0) MarkDuplicates. Reads were filtered with samtools requiring a minimum mapping quality of 20, a maximum fragment length of 2000, including only reads mapped in a proper pair and excluding secondary alignment, duplicate reads and reads failing quality checks (samtools view -F 3840 -f 3 -q 20). Reads where then separated in two bam files belonging to the endogenous species (*D. melanogaster*) and the exogenous species (*D. virilis*) per spike-in CUT&Tag sample. Non-spike-in normalized signal tracks (bigwig) were obtained using bamCoverage (from deeptools^97^) with options “--binSize 10 --effectiveGenomeSize (*D. mel:* 125464728, *D. vir:* 206026697) --extendReads --exactScaling --ignoreDuplicates” for both species. The quality of the spike-in CUT&Tag libraries was extensively checked with an array of tools, as described above for the ChIP-seq. The library complexity was evaluated with Picard EstimateLibraryComplexity (v. 2.7.1.1). The fingerprint (enrichment of signal) was evaluated with deeptools^97^ plotFingerprint. Correlation between samples was checked with deeptools multiBigwigSummary and plotCorrelation^97^ (v. 3.0.2.0) based on 10bp non-spike-in normalized bigwig files, separately on *D. melanogaster* and *D. virilis* bigwigs. QC results were summarized and compared across samples with multiQC^100^ (v. 1.13). Moreover, peak calling in *D. virilis* was used as a quantitative measure of signal enrichment in the exogenous species (following the same IDR peak calling pipeline described above) for Dark and control (*yw)* samples. In particular, *D. melanogaster* and *D. virilis* peak calls show a reasonable concordance in the number of called peaks across species for the same antibody. The spike in Cut&Tag peaks and differential peaks are provided in Table S4.

##### Spike-in normalization

Spike-in normalization factors were obtained by counting the number of reads in the exogenous species bam file. The spike-in factors were then normalized by dividing the counts by the geometric mean of the spike-in factors within the same contrast. The normalization was run independently for each contrast (e.g. one contrast corresponds to *chip:iLEXYs* Chip antibody dark [two replicates] and blue light [two replicates] samples) to ensure samples comparability. Spike-in normalized signal tracks (bigwig) were obtained (for *D. melanogaster* only) using bamCoverage with options “--binSize 10 --effectiveGenomeSize 125464728 --extendReads --exactScaling --ignoreDuplicates --normalizeUsing ‘None’” and option “--scaleFactor” with the value corresponding to the sample normalized spike-in factor. Coverage for replicate was then averaged with bigwigCompare (from deeptools) with options “--operation mean --binSize 10” (Fig. 5B,C). Peaks were called using the IDR pipeline based on peak summits, as described for ChIP-seq above. Alignments were filtered to include only reads with a maximum insert size of 450. After peak calling using macs2^102^ (v. 2.2.7.1), read counts were normalized across samples with the R package DESeq2^114^ using the reverse (DESeq multiplies the counts by sizeFactor while bamCoverage divides the coverage by the scaleFactor) of the above-described normalized spike-in factor as size factor (DESeq2 library normalization) for each sample. Fig. 7E-J reports the average DESeq2 corrected CUT&Tag signal between replicates. Significantly differential signal in control (*chip:iLEXYs* dark or *yw* samples) vs treatment (*chip:iLEXYs* blue light or *zelda^-/-^*) is defined as FDR < 0.05 and a |log2FC| > 0.5 (Fig. 7E-J red dots and Table S4).

#### Spike-in ATAC-seq

##### Mapping, coverage tracks, spike-in normalization, Library quality control

The spike-in ATAC-seq analysis was run using the same pipeline of the spike-in CUT&Tag described above, including the quality controls. The spike-in ATAC-seq dataset included two contrasts (the first contrast corresponds to *chip:iLEXYs* dark [two replicates] and blue light [two replicates] samples; the second to control (*yw)* [two replicates] and *zelda^-/-^* [two replicates] samples). The peak calling was run as described for the spike-in CUT&Tag with the following modification: MACS2 summits were expanded by 100bp upstream and downstream; IDR peaks were expanded by 300bp upstream and downstream to adjust to the more focused signal from ATAC-seq compared to CUT&Tag (Table S6).

#### Hi-C

##### Mapping and contact matrix normalization

Reads were aligned and filtered using the HiCUP pipeline^117^ (v. 0.9.2) using options “--zip --nofill --bowtie2 bowtie2 --threads” with a digested genome based on DpnII restriction sites. Pairs were obtained with bam2pairs from pairix^118^ (v. 0.3.7) only for major chromosomes (chr2L, chr2R, chr3L, chr3R, chr4, chrX, chrY). Pairs were aggregated into a 1kb contact matrix with cooler^119^ (v. 0.8.5) cload pairs with options “-c1 2 -p1 3 -c2 4 -p2 5 chromosome_size.txt:1000”. Matrix were then coarsened to 2kb, 5kb, 10kb and 20kb resolutions with cooler coarsen. Duplicates were merged with cooler merge to increse signal depth. To ensure comparability between replicates, matrices were scaled to the lowest number of contacts with HiCExplorer^120^ (v. 3.5.3) tool hicNormalize using option “--normalize smallest”. Finally, the matrix counts were balanced with cooler balance and options “--ignore-diags 0 --max-iters 10000 --cis-only --force”. Differential signal between dark and blue light samples was obtained with hicCompareMatrices and option “--operation log2ratio”.

##### Library quality control

The quality of the libraries and mapping was checked with HiCUP QC reports, complemented with HiCExplorer QC report. Duplicates were kept separate (skipping the merge cooler merge step) and processed in the same way as described above for quality control. Concordance of counts vs distance across replicates was tested with hicPlotDistVsCounts (from HiCExplorer). Correlation between replicates and across conditions was checked with hicCorrelate. Compartmentalization was quantified with cooltools^121^ (v. 0.4.0) **‘**call-compartment’ and ‘saddle’. The Hi-C signal was extensively checked manually with hicPlotTADs and by comparing it to external experiments^46,93^.

##### Pileups, loops signal quantification and insulation scores

Hi-C pileups were generated with coolpup.py^122^ (v. 0.9.5) with options “--pad 10 --maxdist 100000 --maxshift 100000 --minshift 11000” based on 1kb resolution balanced matrices (Fig. 6A). Loop intesity was then quantified with chromosight quantify^123^ (v. 1.3.3) for dark and blue light matrices (Fig. 6B). Categories of loops were defined as: 1) high-confidence, manually annotated loops^45^, 2) loops where one of the anchors is a promoter bound by Chip and the other is an enhancer bound by Chip (Fig. 6B), 3) loops where both anchors are promoters bound by Chip, 4) loops where both anchors are enhancer bound by Chip, 5) loops where one of the anchors is a promoter of a differentially expressed gene and the other is an enhancer bound by Chip, 6) loops where one of the anchors is a promoter of a differentially expressed gene bound by Chip and the other is an enhancer bound by Chip, 7) loops where both anchors are Chip peaks. Throughout these analyses, Chip peaks refer to Chip ChIP-seq peaks at 2-4h, and differentially expressed genes refer to differentially expressed genes in dark vs blue light at 2-4h.

Finally, insulation scores where obtained with hicFindTADs and options “-- correctForMultipleTesting fdr --thresholdComparisons 0.01 --delta 0.01 --step 5000 --minDepth 15000 --maxDepth 200000” on 5kb resolution balanced matrices. We then compared the distribution of insulation scores on annotated TAD boundaries called in^46^ (Fig. 6D).

#### Capture-C

##### Capture-C probe design

To investigate the role of Chip (Lbd1) and Zelda in enhancer-promoter interactions, we designed 501 Capture-C probes targeting different categories of elements, as follows:

1. 72 probes targeting the promoters of genes differentially expressed in both *zelda^-/-^*and *chip:iLEXYs* blue light and that are bound both by Chip (this study, ChIP-seq peaks 2-4h) and Zelda (ChIP-seq 2-2.5h and 3-3.5h peaks from^40^).
2. 112 probes targeting a characterised enhancer (>5kb away from any known TSS and overlapping an active characterised enhancer (from CAD4^36^)) bound both by Chip (this study, ChIP-seq peaks 2-4h) and Zelda (ChIP-seq 2-2.5h and 3-3.5h peaks from^40^).
3. 90 probes targeting enhancers bound by Chip and not bound by Zelda.
4. 107 probes targeting enhancers bound by Zelda and not bound by Chip.
5. 25 probes targeting the promoters of genes essential for early *Drosophila* development that were not selected following other criteria (e.g. *eve*, *en*, *hb*, *inv*…)
6. 30 probes targeting control promoters selected as: non-differentially expressed in *zelda^-/-^*and *chip:iLEXYs* blue light, expressed in *chip:iLEXYs* dark, not bound by both Chip (this study, ChIP-seq peaks 2-4h) and Zelda (ChIP-seq 2-2.5h and 3-3.5h peaks from^40^).
7. 65 probes targeting control enhancers selected as: TSS-Distal (>2kb from any known TSS), ATAC-seq peaks (this study, ATAC-seq *chip:iLEXYs* dark), not bound by both Chip and Zelda.

The final list was manually filtered after sorting the elements for the intensity of the signal of the target transcription factor(s) (Table S5).

The probes were designed using the python package oligo (v. 0.1.2) with options “Capture -g dm6 -o 120 -e DpnII” and STAR^124^ for off-targets identification.

Mapping, coverage tracks and differential analysis considering fragments with a distance more than 2kb or less than 100kb from the bait was performed as described in^47^.

